# Moderating carbohydrate digestion rate promotes metabolic flexibility in mice

**DOI:** 10.1101/2023.01.08.523060

**Authors:** Anna M.R. Hayes, Clay Swackhamer, Roberto Quezada-Calvillo, Nancy F. Butte, Erwin E. Sterchi, Buford L. Nichols, Bruce R. Hamaker

## Abstract

Superior metabolic flexibility, or the ability to efficiently switch between oxidation of carbohydrate and fat, is inversely associated with obesity and type 2 diabetes. This study examined the impact of dietary carbohydrate digestion rate on metabolic substrate utilization and metabolic flexibility. We employed percent relative cumulative frequency (PRCF) analyses coupled with a new application of modeling using the Mixed Weibull Cumulative Distribution function to examine respiratory exchange ratio (RER) data from wild-type mice and mice lacking the mucosal maltase-glucoamylase enzyme (Mgam, null) under different dietary carbohydrate conditions. We further devised a Metabolic Flexibility Factor (MFF) to quantitate metabolic flexibility, with higher MFF indicating higher metabolic flexibility. The collective results indicated that a diet high in slowly digestible starch exhibited higher metabolic flexibility (MFF) than diets high in resistant starch, sucrose, or fat. These findings show a new-found benefit of consuming slowly digestible carbohydrates for improved metabolic health.

## Introduction

As the prevalence of obesity and type 2 diabetes continues to pose immense burdens worldwide, efforts to prevent or decrease their severity remain crucial in order to sustain a healthy human population. From a metabolic standpoint, research from recent decades has revealed a relationship between mitochondrial metabolic function and insulin resistance occurring with type 2 diabetes and obesity ^1-4^. Specifically, alterations in the ability of mitochondria to switch between utilization of fat or carbohydrate according to nutritive conditions for oxidation, a concept termed “metabolic flexibility”, may precede insulin resistance, contributing to the development of type 2 diabetes and obesity ^4,5^. Having poor metabolic flexibility (i.e., metabolic inflexibility) is thought to be rooted in a rigidity in mitochondrial substrate selection, such that mitochondria have hindered ability to freely switch fuel choice according to nutritional circumstances ^6^. Metabolic flexibility has been studied extensively in the context of disease states ^2,7,8^, exercise ^3,9^, and fasting ^2,10^. Although substrates used for metabolism are largely provided through the diet, a relatively small number of studies have focused on the connection between diet and metabolic flexibility ^11-13^. In particular, there is a need to better understand the role of carbohydrate digestibility on metabolic flexibility due to their major function as a metabolic fuel substrate ^14,15^.

Dietary carbohydrates constitute the main source of energy in most diets ^16^, and thus determining how carbohydrates can be used to help prevent or reduce the detrimental side effects of obesity and type 2 diabetes could be key for combating these public health challenges, and ultimately to enable improved food production and processing efforts to sustain a healthier planet. A challenge is to better define and understand carbohydrate ‘quality’. Basic differences in the types of carbohydrates – such as sugar (sucrose), starch, and fiber – are widely recognized. Yet, within these types, not all sugars, starches, or fibers are the same. For starches, differences in botanical source ^17,18^; macro-, micro-, or fine structure ^19-22^; food form ^23,24^; processing and preparation ^25-27^; and other factors give rise to different post-ingestive (acute) effects on the body that lead to other impacts in the long term ^28-30^. These consequences mainly relate to the rate at which carbohydrate macromolecules are hydrolyzed into simple sugars (digestion rate) as well as the location where such hydrolysis takes place along the small intestine (digestion location) ^31-33^.

The digestion or hydrolysis of starch and other digestible α-glucans to absorbable free glucose is termed α-glucogenesis ^34^, involving six starch-degrading enzymes: salivary α-amylase, pancreatic α-amylase, and the α-glucosidases comprising maltase-glucoamylase (Mgam; N-terminal and C-terminal) and sucrase-isomaltase (Si; N-terminal and C-terminal), each with two enzyme activities. These six enzymes complement each other and may be adaptive to evolutionary and cultural variations in food and feeding/eating. Substantial research has been conducted to better understand the catalytic sites, activities, and substrate specificities of these enzymes from humans and other animals ^34-43^. However, it is incompletely understood how α-glucogenesis related to carbohydrate digestion rate may impact metabolism and energy partitioning. Most of the *in vivo* work on slowly digestible starch (SDS) to date has focused on glycemic response or physiological/hormonal responses, and the work on resistant starch (RS) and fiber has generally focused on glycemic response, fermentation, and the gut microbiome. SDS is hydrolyzed slowly in the small intestine and possesses a measured and sustained blood glucose release, while rapidly digestible starch (RDS) is hydrolyzed rapidly and leads to a pronounced postprandial blood glucose spike ^44-47^. Furthermore, RS passes into the large intestine to undergo fermentation, resulting in the production of short chain fatty acids (SCFAs) ^44,48^. In terms of metabolism, aside from the study by Fernández-Calleja et al. ^14^ showing a more pronounced carbohydrate oxidation response in female mice compared to male mice fed a high-amylose diet (diet with 60% of carbohydrate component from amylose) and the study by Salto et al. ^15^ indicating increased fat oxidation during catch-up growth of mice fed a SDS diet, to our knowledge no reports exist regarding how the amount or type of SDS or RS affects metabolism and partitioning of carbohydrate vs. fat for energy. Furthermore, no studies on SDS and metabolic flexibility have been done. Sophisticated tools have been developed that allow for simultaneous, continuous, and prolonged measurement of respiratory exchange ratio (RER), ^13^CO_2_, and fermentation indicators (i.e., H_2_ and CH_4_) using an enhanced indirect calorimetry system ^14,49-51^, which makes possible investigations of carbohydrate partitioning for oxidation as well as metabolic flexibility.

In the present study, we investigated metabolic fuel utilization (RER) and metabolic flexibility of diets containing carbohydrates with different digestibilities (Figure 1), comprised either of materials themselves differing in digestibility, or decreasing carbohydrate digestion rate through knock-out of Mgam or increasing carbohydrate digestion rate through addition of an external starch-degrading enzyme. The Mgam null mouse exhibits a 40% reduction in mucosal digestion of glucogenesis, but normal energy expenditure ^34,36,37^. For diets, we selected a starch that is slowly digestible (raw corn starch, RCS) ^20,21,46^ and a starch that is mostly resistant to digestion in the small intestine (high-amylose starch, HAS) ^52,53^. We also included a diet that contained sucrose, a readily available carbohydrate ^54^, and a diet high in fat (low in carbohydrate) as comparators. Thus, the six different diets were as follows: High SDS (53% RCS); Intermediate SDS (35% RCS + 18% HAS); Intermediate RS (18% RCS + 35% HAS); High RS (53% HAS); Sucrose (65% sucrose); and High-fat (42% fat). Our initial findings showed that mice consuming the four types of SDS and RS diets did not exhibit differences in energy expenditure (Figure S1). These findings were intriguing because the extent of carbohydrate digestibility differed markedly among the diets, and thus suggest that the mice metabolically adapted and therefore underwent a change in metabolic fuel utilization.

**Figure 1.**
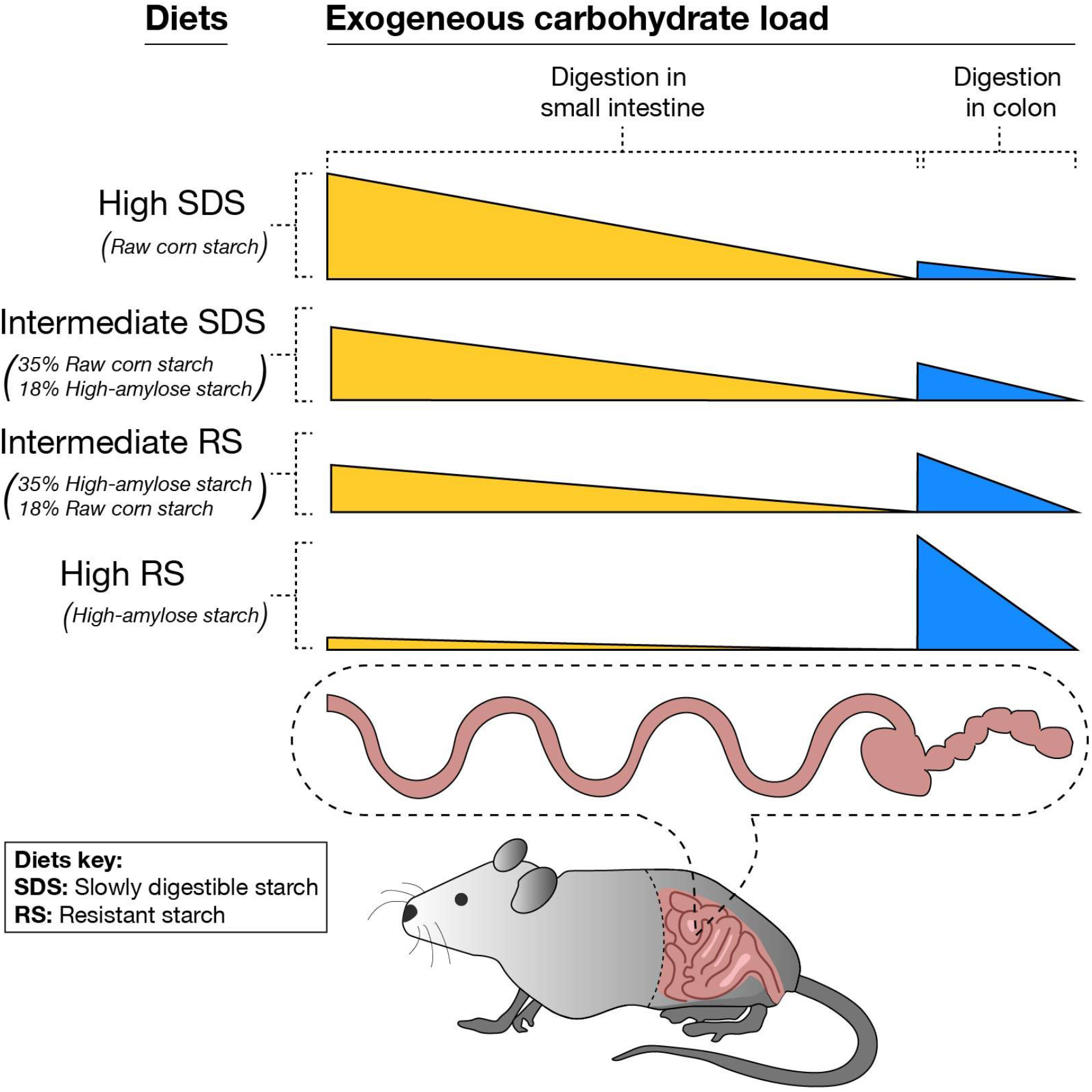
Digestion profiles of experimental carbohydrate diets. Slowly digestible starch (SDS, represented in yellow) is hydrolyzed in the small intestine whereas resistant starch (RS, represented in blue) is fermented in the colon. Total carbohydrate content of each diet was held constant at 53%.

We hypothesized that carbohydrates with different digestibilities would affect the utilization of glucose vs. fat as substrates for energy. Additionally, we reasoned that dietary carbohydrates with slow digestion rates would decrease the metabolic gridlock ^6^ that can occur due to the rapid influx of glucose into mitochondria following consumption of a rapidly digestible carbohydrate, and thus result in enhanced metabolic flexibility. To thoroughly probe this hypothesis, we altered carbohydrate digestibility/digestion 1) in diets with different proportions of slowly digestible starch and resistant starch, 2) using Mgam knockout (null) mice to reduce starch digestibility, and 3) adding amyloglucosidase (AMG) to increase digestibility (see experimental design in Figure S2). The impacts of these alterations were assessed by measuring respiratory exchange ratio (RER), the ratio of carbohydrate oxidation to fat oxidation, and metabolic flexibility in mice.

The measurement of RER, which represents the ratio of carbohydrate oxidation to fat oxidation, was the primary study outcome, as measured by indirect calorimetry. Because the existing approaches for analyzing RER provided limited insight into substrate partitioning and metabolic flexibility, we developed two new approaches for examining specific characteristics of substrate oxidation and metabolic flexibility through RER:

1. Employing percent relative cumulative frequency (PRCF) analysis as proposed by Riachi et al. ^55^, plus adding a new modeling approach with the Mixed Weibull Cumulative Distribution function to differentiate carbohydrate vs. fat oxidation.
2. Using this approach, we constructed an intuitive numerical factor, which we termed the Metabolic Flexibility Factor (MFF), to quantify the proportion of RER values falling within “ideal” ranges of RER for fat and carbohydrate oxidation that capture a metabolically sensitive and flexible state.

The application of such approaches revealed how different digestive rates of carbohydrate-based diets or dietary conditions have distinct effects on metabolic substrate utilization.

## Results

### Mean respiratory exchange ratio (RER) per 24 h

Three-way ANOVA conducted on the mean respiratory exchange ratio (RER) values per 24 h cycle for the groups (diet × genotype × cycle [without/with AMG]) revealed main effects for diet and cycle (*p*<0.0001 for both). The interaction term for diet × genotype × cycle was also significant (*p*=0.0001), so *post hoc* comparisons were made among the 24 groups (Table 1). The Sucrose diet had the highest mean RER values, with null mice (both with and without AMG) having significantly higher RER than all other groups and wild-type mice having significantly higher RER than all other groups except the wild-type mice fed the High RS diet when given AMG and the null mice fed the Sucrose diet. This suggests that the Sucrose diet yielded a greater extent of carbohydrate oxidation. The relatively high amount of RDS contributed by Novelose 260 may have given rise to the high mean RER for the High RS diet (see Table S2), and the relatively high amount of SDS contributed by raw corn starch may have resulted in the low mean RER for the High SDS diet. AMG supplementation also had a clear effect for both genotypes fed the High SDS diet and the null mice fed the High RS diet. We then sought to take a deeper look at these findings and examine metabolic flexibility.

**Table 1.**
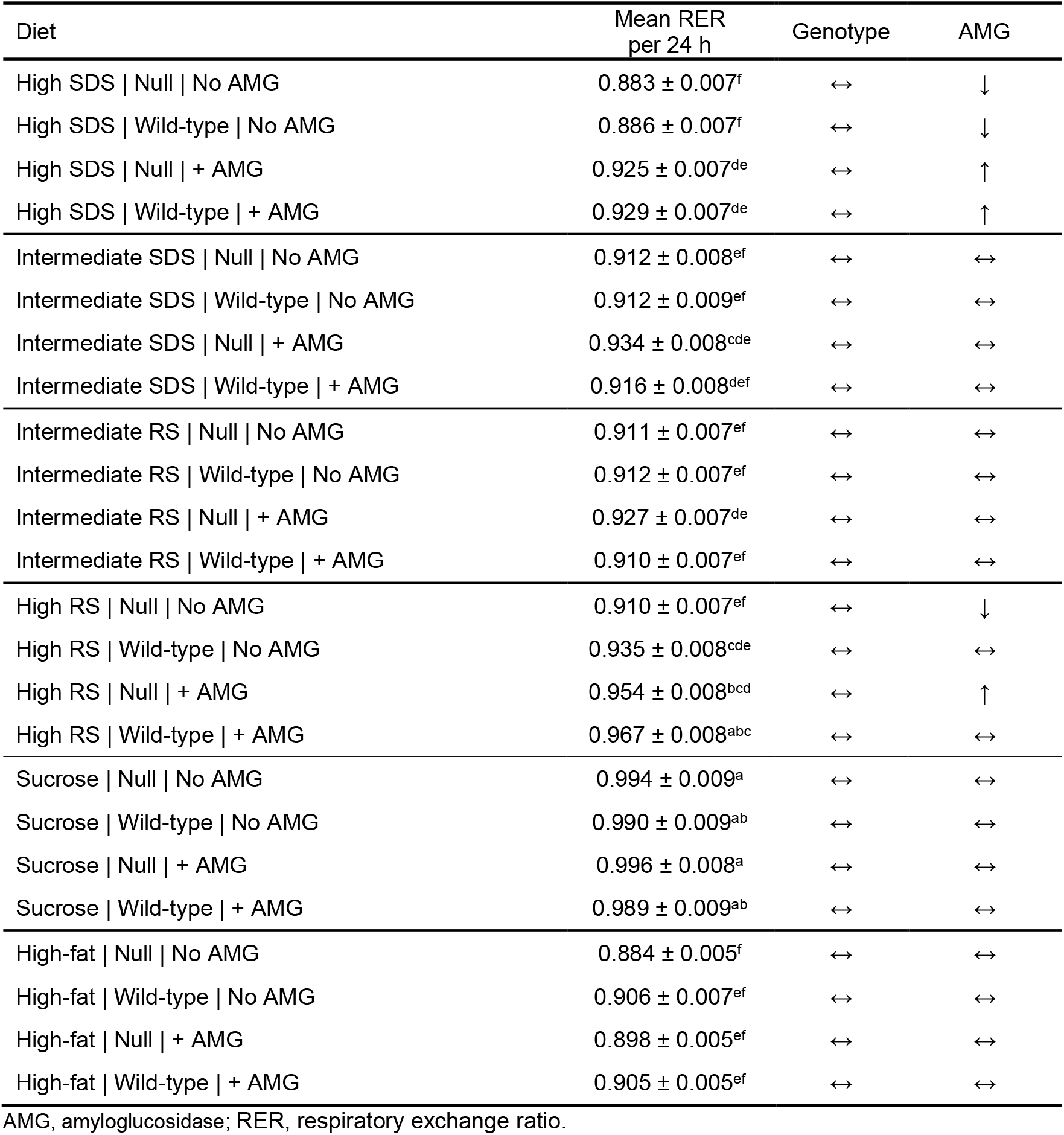
Mean respiratory exchange ratio (RER) per 24 h for each group (diet × genotype × cycle [without/with amyloglucosidase, AMG]). Mean values shown with ± standard error of the mean. Values not sharing the same letter are significantly different (*p*<0.05; diet × genotype × cycle was significant [*p*=0.0001], so *post hoc* comparisons were made among the 24 groups). Arrows in columns for genotype and AMG breakdown the effects of these two factors *within* each diet.

| Diet | Mean RER<br>per 24 h | Genotype | AMG |
| --- | --- | --- | --- |
| High SDS \| Null \| No AMG | 0.883 ± 0.007<sup>f</sup> | ↔ | ↓ |
| High SDS \| Wild-type \| No AMG | 0.886 ± 0.007<sup>f</sup> | ↔ | ↓ |
| High SDS \| Null \| + AMG | 0.925 ± 0.007<sup>de</sup> | ↔ | ↑ |
| High SDS \| Wild-type \| + AMG | 0.929 ± 0.007<sup>de</sup> | ↔ | ↑ |
| Intermediate SDS \| Null \| No AMG | 0.912 ± 0.008<sup>ef</sup> | ↔ | ↔ |
| Intermediate SDS \| Wild-type \| No AMG | 0.912 ± 0.009<sup>ef</sup> | ↔ | ↔ |
| Intermediate SDS \| Null \| + AMG | 0.934 ± 0.008<sup>cde</sup> | ↔ | ↔ |
| Intermediate SDS \| Wild-type \| + AMG | 0.916 ± 0.008<sup>def</sup> | ↔ | ↔ |
| Intermediate RS \| Null \| No AMG | 0.911 ± 0.007<sup>ef</sup> | ↔ | ↔ |
| Intermediate RS \| Wild-type \| No AMG | 0.912 ± 0.007<sup>ef</sup> | ↔ | ↔ |
| Intermediate RS \| Null \| + AMG | 0.927 ± 0.007<sup>de</sup> | ↔ | ↔ |
| Intermediate RS \| Wild-type \| + AMG | 0.910 ± 0.007<sup>ef</sup> | ↔ | ↔ |
| High RS \| Null \| No AMG | 0.910 ± 0.007<sup>ef</sup> | ↔ | ↓ |
| High RS \| Wild-type \| No AMG | 0.935 ± 0.008<sup>cde</sup> | ↔ | ↔ |
| High RS \| Null \| + AMG | 0.954 ± 0.008<sup>bcd</sup> | ↔ | ↑ |
| High RS \| Wild-type \| + AMG | 0.967 ± 0.008<sup>abc</sup> | ↔ | ↔ |
| Sucrose \| Null \| No AMG | 0.994 ± 0.009<sup>a</sup> | ↔ | ↔ |
| Sucrose \| Wild-type \| No AMG | 0.990 ± 0.009<sup>ab</sup> | ↔ | ↔ |
| Sucrose \| Null \| + AMG | 0.996 ± 0.008<sup>a</sup> | ↔ | ↔ |
| Sucrose \| Wild-type \| + AMG | 0.989 ± 0.009<sup>ab</sup> | ↔ | ↔ |
| High-fat \| Null \| No AMG | 0.884 ± 0.005<sup>f</sup> | ↔ | ↔ |
| High-fat \| Wild-type \| No AMG | 0.906 ± 0.007<sup>ef</sup> | ↔ | ↔ |
| High-fat \| Null \| + AMG | 0.898 ± 0.005<sup>ef</sup> | ↔ | ↔ |
| High-fat \| Wild-type \| + AMG | 0.905 ± 0.005<sup>ef</sup> | ↔ | ↔ |

AMG, amyloglucosidase; RER, respiratory exchange ratio.

### Novel strategy to examine metabolic fuel selection

Percent relative cumulative frequency was initially proposed as a means to analyze RER data by Riachi et al. ^55^. Because this approach involves sorting the data from lowest to highest, the temporal nature of the dataset is lost. In exchange, it describes the distribution of RER values, which allows for detailed examination of the overall clustering (or lack thereof) of RER values observed. PRCF of RER was analyzed per individual mouse and, to take this approach one step further, fit to the Mixed Weibull Cumulative Distribution function (Eq.1; Figure 2), with constraints applied as described in the Methods section (notably, the Eq. 2 gamma function).

**Figure 2.**
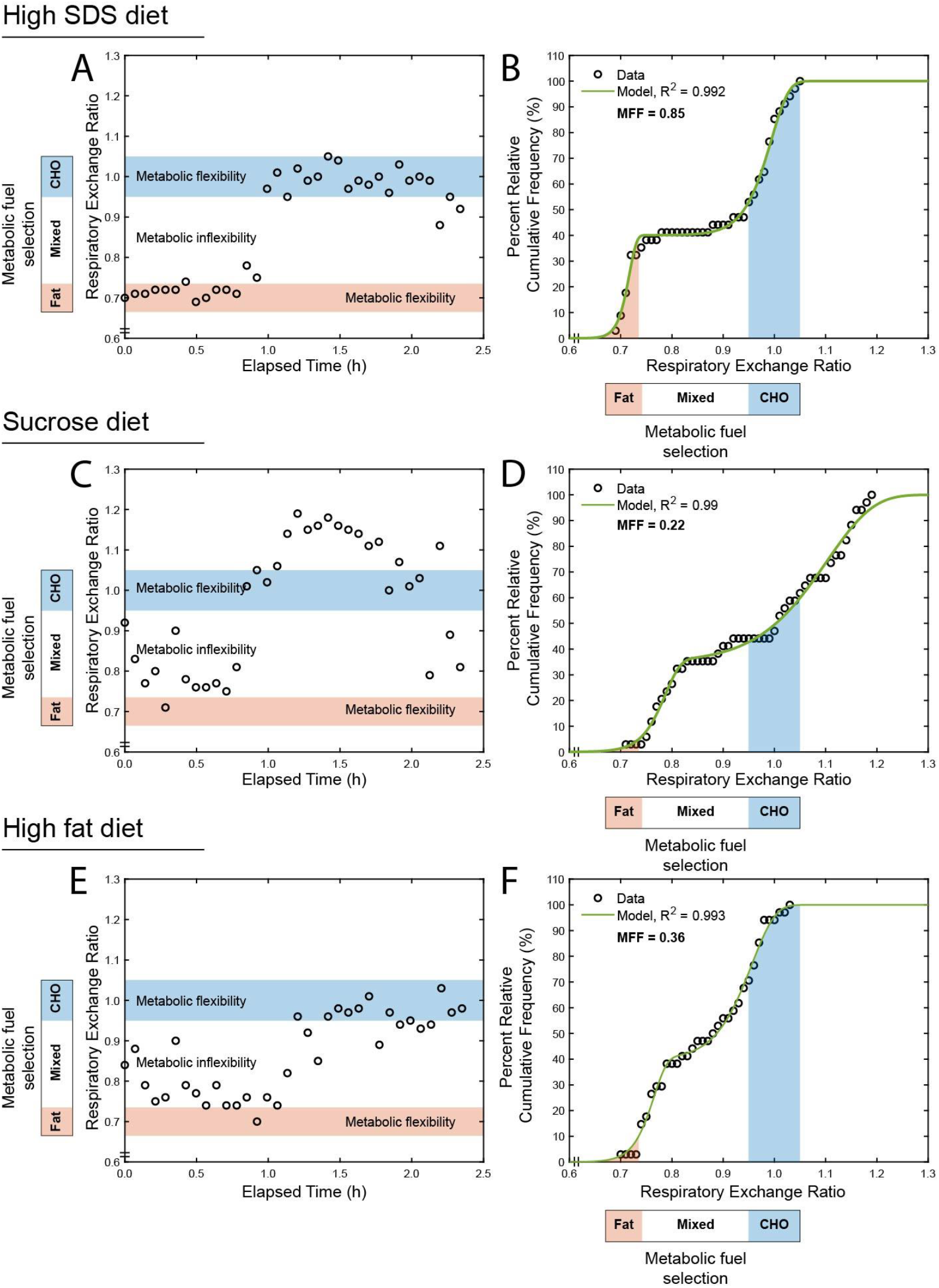
Representative respiratory exchange ratio (RER) curves from individual mice on the High SDS diet (A), the Sucrose diet (C), the High fat diet (E), and their corresponding percent relative cumulative frequency (PRCF) curves (B, D, and F, respectively).

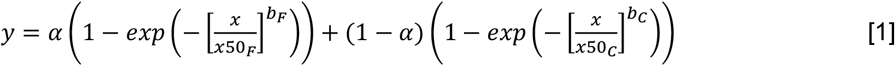

Where:

*y* = percent relative cumulative frequency (PRCF; 0 to 100%);

*α* = mixing weight parameter that represents the proportion of the first mode;

*x*50_*F*_ = median respiratory exchange ratio (median RER) for the <u>first</u> mode;

*b*_*F*_ = distribution breadth constant for <u>first</u> mode (dimensionless), indicative of slope for the first mode;

*x*50_*C*_ = median respiratory exchange ratio (median RER) for the <u>second</u> mode;

*b*_*C*_ = distribution breadth constant for the <u>second</u> mode (dimensionless), indicative of slope for the second mode.

All mean parameter estimates from the Mixed Weibull Cumulative Distribution are shown in Table 2. To aid in interpretation of these parameters, representations of each with theoretical curves are shown in Figure 3. In the subpanels of this figure, one parameter is altered while all others are held constant to visualize how differences in distribution of the data are reflected in the appearance of the Mixed Weibull Cumulative Distribution function.

**Table 2.**
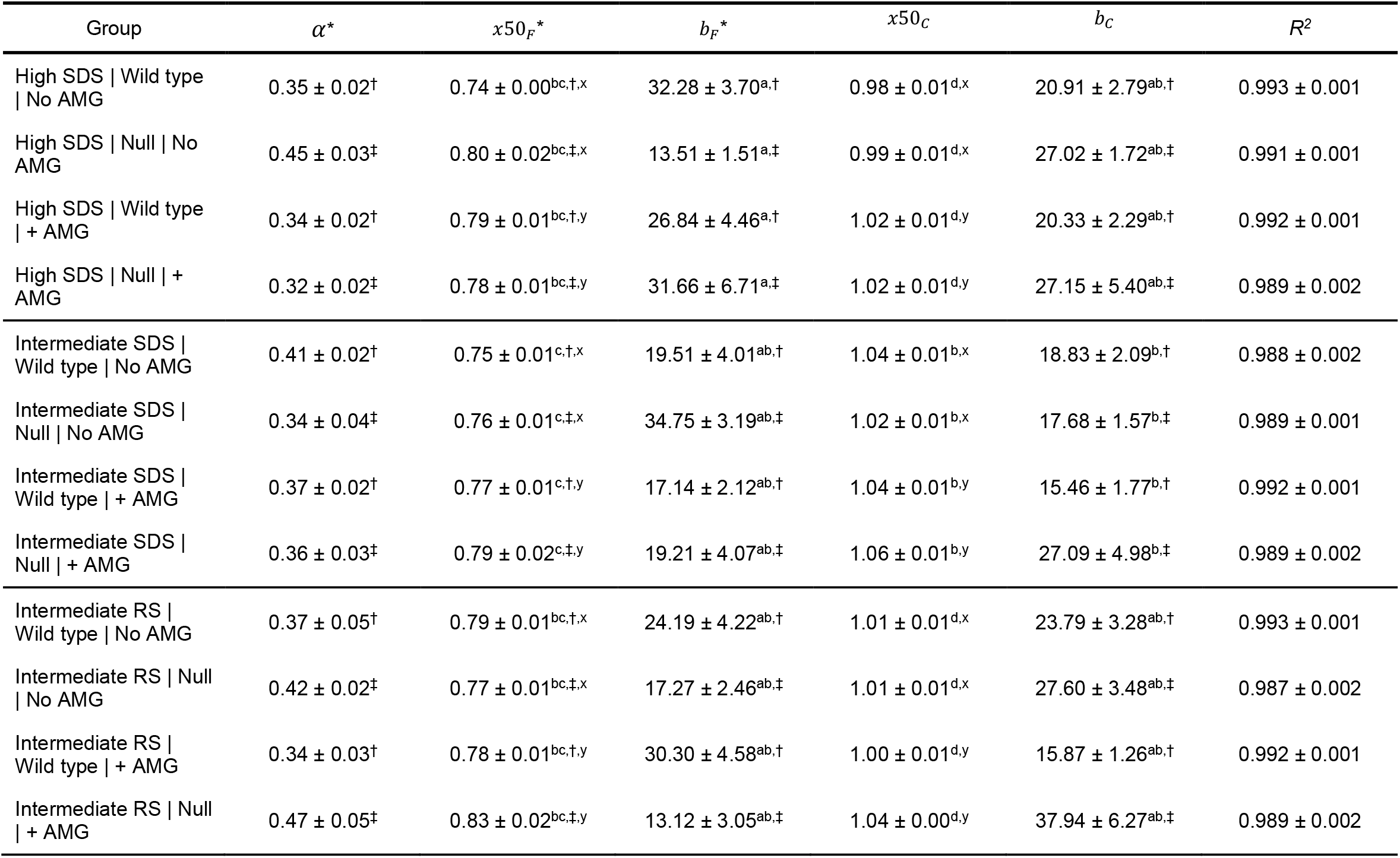
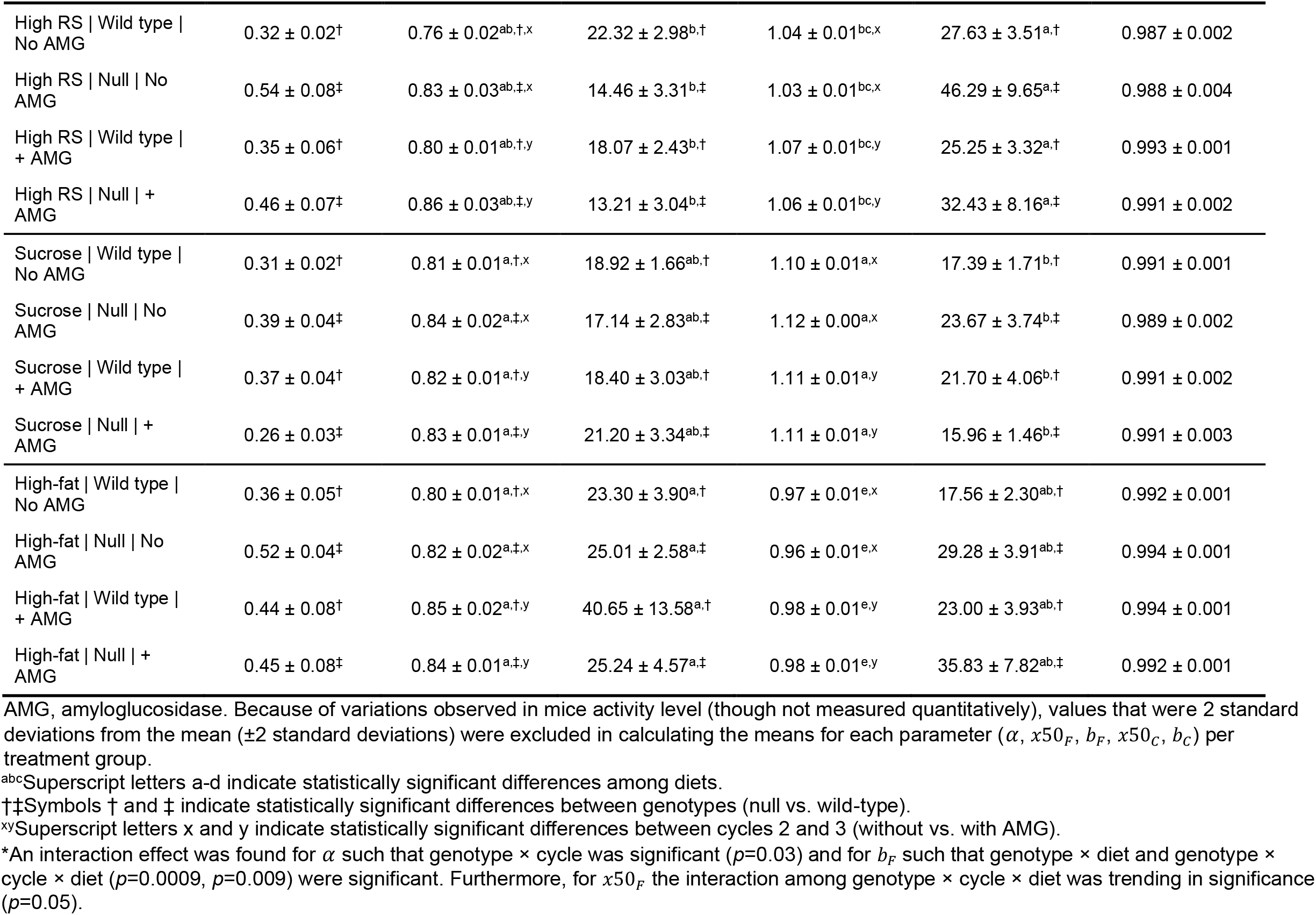
Mean Mixed Weibull Cumulative Distribution parameter estimates (*α, x*50_*F*_, *b*_*F*_, *x*50_*C*_, *b*_*C*_) for respiratory exchange ratio (RER) from individual percent relative cumulative frequency (PRCF) analyses for individual mice. *R*^*2*^ is also shown as an indicator of goodness of fit. Values are means ± standard error of the mean. Groups not sharing the same letters or symbols are significantly different (*p*<0.05).

**Figure 3.**
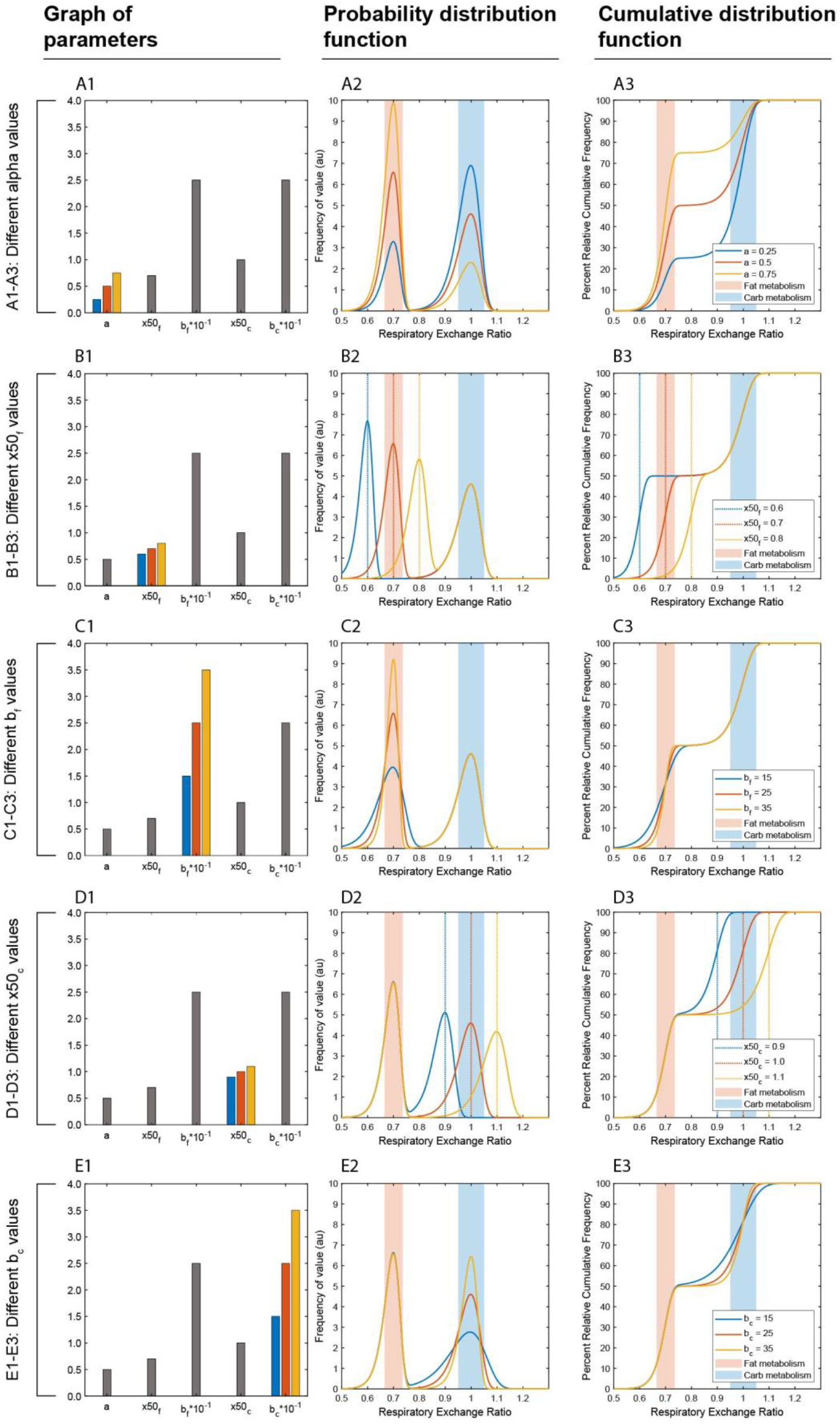
Mixed Weibull parameter exploration for percent relative cumulative frequency (PRCF) analysis of respiratory exchange ratio (RER). Note that all values shown in this figure are theoretical in order to demonstrate the meaning of the parameters. Shaded areas indicate ideal RER ranges for fat oxidation (red) and carbohydrate oxidation (blue). **<u>A1-A3:</u>** Different *α* but the same *x*50_*F*_, *x*50_*C*_, *b*_*F*_, and *b*_*C*_. Bar graph of all parameters, with *α* indicated in color (A1). Probability distribution function indicating theoretical distributions of data with three different *α* values (A2). Cumulative distribution function [Mixed Weibull Cumulative Distribution function] for the same *α* values (A3). Smooth curves indicate curves of distribution. Related to RER, the first mode represents fat oxidation and the second mode represents carbohydrate oxidation. As shown in the figure, greater *α* causes a greater proportion of the distribution to be allocated to the first mode and consequently a lesser proportion of the distribution constitutes the second mode. In practical terms, this means a greater *α* shifts RER toward increased fat oxidation. **<u>B1-B3:</u>** Different *x*50_*F*_ but the same *α, x*50_*C*_, *b*_*F*_, and *b*_*C*_. Bar graph of all parameters, with *x*50_*F*_ indicated in color (B1). Probability distribution function indicating theoretical distributions of data with three different *x*50_*F*_ values (B2). Cumulative distribution function [Mixed Weibull Cumulative Distribution function] for the same *x*50_*F*_ values (B3). Smooth curves indicate curves of distribution. Vertical colored lines indicate *x*50_*F*_ RER values, which are the median RER values of the carbohydrate mode of the respective distributions. Related to RER, the first mode represents fat oxidation and the second mode represents carbohydrate oxidation. As shown in the figure, greater *x*50_*F*_ shifts the curve representing the first mode to the right, which signifies a higher median RER value in the fat oxidation mode. The further *x*50_*F*_ differs from 0.70, the *less* fat oxidation. **<u>C1-C3:</u>** Different *b*_*F*_ but the same *α, x*50_*F*_, *x*50_*C*_, and *b*_*C*_. Bar graph of all parameters, with *b*_*F*_ indicated in color (C1). Probability distribution function indicating theoretical distributions of data with three different *b*_*F*_ values (C2). Cumulative distribution function [Mixed Weibull Cumulative Distribution function] for the same *b*_*F*_ values (C3). Smooth curves indicate curves of distribution. Related to RER, the first mode represents fat oxidation and the second mode represents carbohydrate oxidation. As shown in the figure, greater *b*_*F*_ steepens the curve representing the first mode, which signifies a smaller spread of RER values in the fat oxidation mode. We interpret this to suggest more efficient switching to fat oxidation and thus enhanced metabolic flexibility. **<u>D1-D3:</u>** Different *x*50_*C*_ but the same *α, x*50_*F*_, *b*_*F*_, and *b*_*C*_. Bar graph of all parameters, with *x*50_*C*_ indicated in color (D1). Probability distribution function indicating theoretical distributions of data with three different *x*50_*C*_ values (D2). Cumulative distribution function [Mixed Weibull Cumulative Distribution function] for the same *x*50_*C*_ values (D3). Smooth curves indicate curves of distribution. Vertical colored lines indicate *x*50_*C*_ RER values, which are the median RER values of the carbohydrate mode of the respective distributions. Related to RER, the first mode represents fat oxidation and the second mode represents carbohydrate oxidation. As shown in the figure, greater *x*50_*C*_ shifts the curve representing the second mode to the right, which signifies a higher median RER value in the carbohydrate oxidation mode. The further *x*50_*C*_ differs from 1.00, the less carbohydrate oxidation. **<u>E1-E3:</u>** Different *b*_*C*_ but the same *α, x*50_*F*_, *x*50_*C*_, and *b*_*F*_. Bar graph of all parameters, with *b*_*C*_ indicated in color (E1). Probability distribution function indicating theoretical distributions of data with three different *b*_*C*_ values (E2). Cumulative distribution function [Mixed Weibull Cumulative Distribution function] for the same *b*_*C*_ values (E3). Smooth curves indicate curves of distribution. Related to RER, the first mode represents fat oxidation and the second mode represents carbohydrate oxidation. As shown in the figure, greater *b*_*C*_ steepens the curve representing the second mode, which signifies a smaller spread of RER values in the carbohydrate oxidation mode. We interpret this to suggest more efficient switching to carbohydrate oxidation and thus enhanced metabolic flexibility. au, arbitrary unit; PRCF, percent relative cumulative frequency; RER, respiratory exchange ratio.

The PRCF curves of the RER data from this study generally appeared bimodal, with one mode ranging from approximately 0.65 to 0.85 RER and the other mode ranging from approximately 0.95 to 1.03 RER (Figure 4). Because an RER of 0.70 indicates fat is being used as the predominant fuel source, and an RER of 1.00 indicates carbohydrate is being used as the predominant fuel source, we propose that the two modes in our PRCF curves represent fat oxidation and carbohydrate oxidation, respectively. Using this interpretation, the *x*50_*F*_ value from modeling with the Mixed Weibull Cumulative Distribution function represents the median RER of the fat oxidation mode, and the *x*50_*C*_ value represents the median RER of the carbohydrate oxidation mode. Furthermore, the *b*_*F*_ and *b*_*C*_ values describe the slopes of the fat oxidation and carbohydrate oxidation modes, respectively, on the graphs of PRCF vs. RER (Figures 3-4).

**Figure 4.**
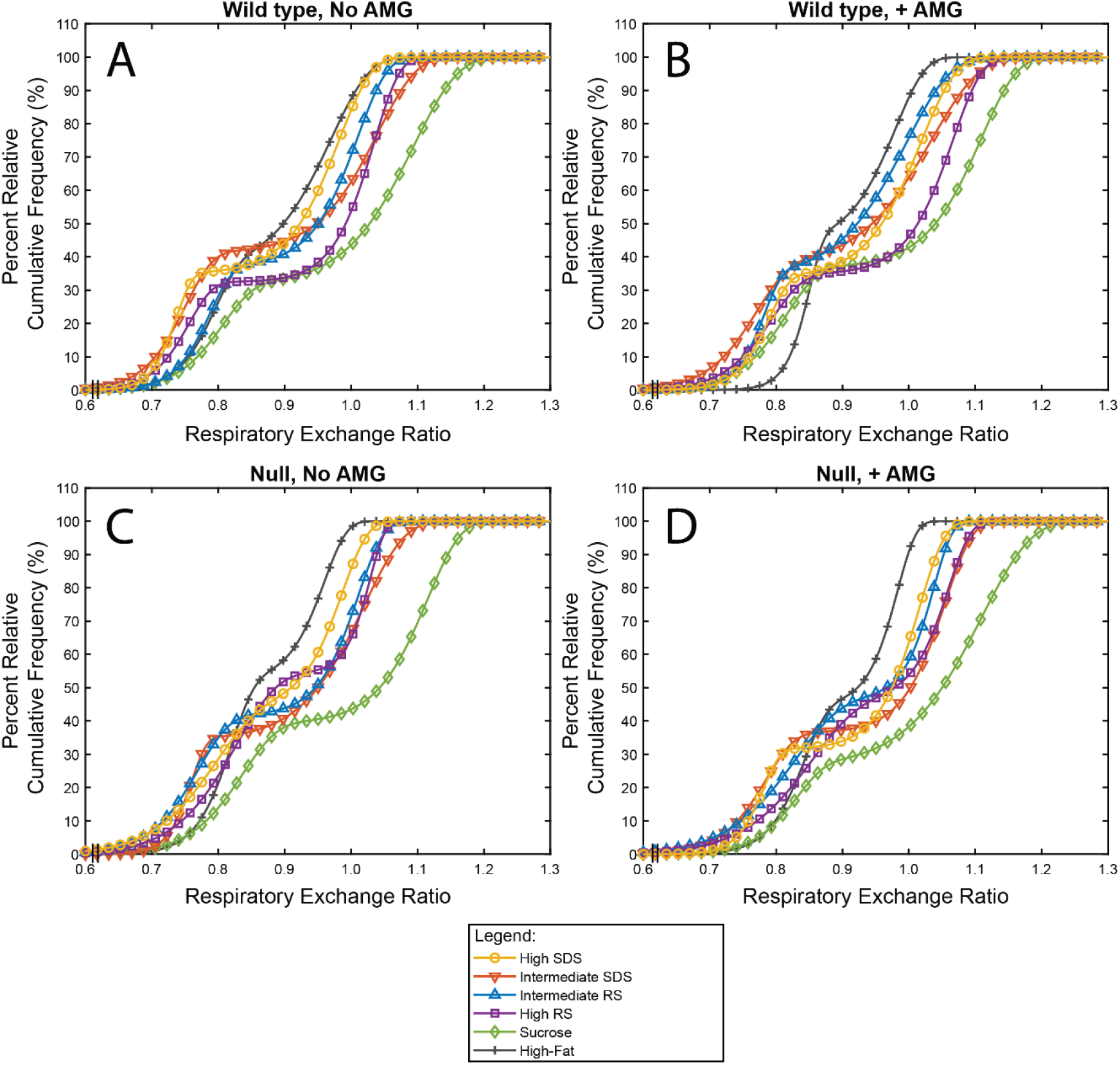
Percent relative cumulative frequency (PRCF) curves of respiratory exchange ratio (RER) modeled using the Mixed Weibull Cumulative Distribution for individual mice and plotted by taking the means of the modeled parameter values. Split by genotype and amyloglucosidase (AMG) supplementation: wild-type without AMG (A), wild-type with AMG (B), null without AMG (C), null with AMG (D). AMG, amyloglucosidase; PRCF, percent relative cumulative distribution; RCS, RER, respiratory exchange ratio.

### Slow carbohydrate digestion promotes fat oxidation

Using the Mixed Weibull Cumulative Distribution approach, three-way ANOVA indicated a main effect of diet for the *x*50_*F*_, *b*_*F*_, *x*50_*C*_, and *b*_*C*_ parameters (*p*<0.0001, *p*=0.0005, *p*<0.0001, and *p*=0.0003, respectively; Table 2, Figure 3, Figure S3).

*Post-hoc* analyses revealed that the High SDS, Intermediate SDS, and Intermediate RS diets had significantly lower *x*50_*F*_ values than the Sucrose and High-fat diets, indicating that they had lower median RER values for the first mode, representing higher fat oxidation, compared to the Sucrose and High-fat diets. Furthermore, the Intermediate SDS diet had lower *x*50_*F*_ than the High RS diet. A lower RER value for the fat oxidation mode suggests that, during periods of fat oxidation, little concurrent oxidation of carbohydrate occurred, resulting in RER values close to the theoretical value that would result from purely fat oxidation (0.7). Notably, the High SDS, Intermediate SDS, and Intermediate RS diets contained greater amounts of RCS, which has a pronounced slow digestion property ^46^. Oxidation of fat within the body is generally driven by the lack of carbohydrate consumed (i.e., displacement of carbohydrate consumption) ^56^. In our study, total carbohydrate content was matched among the different diets, but carbohydrate digestibility varied; this enabled changes in RER to be attributed to carbohydrate digestibility rather than total carbohydrate amount. RER is also affected by body weight, body composition, and level of activity ^56-58^. In our case, the mice were similar in body weight (see the Body Weight section in Supplementary Information) and body composition (data not shown), but we did not measure activity. Future research incorporating activity monitoring would help describe any potential interplay between activity levels, carbohydrate digestion, and fat oxidation. The High-fat diet had higher *x*50_*F*_ values than anticipated (indicating lower fat oxidation), but it is important to note that this diet also contained a relatively high amount of sucrose (Table S1). Additionally, a higher *x*50_*F*_ for this diet could be hallmark of metabolic inflexibility. Specific to the null mice, a previous study showed enhanced activity of sucrase-isomaltase (Si) as partial compensation for the lack of Mgam ^36^, and that Si may have a preferential ability to hydrolyze sucrose and thereby affect RER.

For *b*_*F*_, the High RS diet had significantly lower values than the Sucrose and High SDS diet groups, but all other diets did not differ (Table 2). This finding suggests there was a wider spread in RER values for the first mode (representing fat oxidation) for the High RS diet than for the Sucrose and High SDS diets (Figure 3). A key concept in metabolic flexibility is how substrate utilization changes during the transition from fed to fasted states ^4^. Although the nature of this analysis eliminates time order, broadening in the clusters of RER values in either the fat or carbohydrate oxidation modes could be indicative of a gradual and less complete change in substrate utilization – as opposed to a swift and more complete change, as would be characteristic of a narrower range of values in the mode.

For *x*50_*C*_, representing the median RER of the second mode, diets ranked as follows (highest to lowest *x*50_*C*_): Sucrose diet > High RS diet = Intermediate SDS diet > Intermediate RS diet = High SDS diet > High-fat diet (Table 2). Because *x*50_*C*_ describes carbohydrate oxidation, these findings indicate that greater carbohydrate oxidation was observed for all the carbohydrate-predominant diets (i.e., the High SDS, Intermediate SDS, Intermediate RS, High RS, and Sucrose diets) compared to the High-fat diet. Specifically, the Sucrose diet contained a large portion of readily available carbohydrate (note that the different absorption pathway for fructose ^54^ contributes to carbohydrate oxidation), and thus it had the highest carbohydrate oxidation with mean values of 1.10-1.12. Because *de novo* lipogenesis may increase RER ^59,60^, such high values suggest carbohydrate was being oxidized and stored as fat for the Sucrose diet feeding. As HAS is partly digestible, the High RS diet likely had a portion of starch that was rapidly digestible to drive carbohydrate oxidation (supported by the digestibility data in Table S2). The reason the Intermediate SDS diet had similar mean carbohydrate oxidation to the High RS diet is unclear, but it may also be that the Intermediate RS diet has an optimized ratio of RS to SDS to exert an effect on RER. The importance of the effect size of these differences (values from 1.03-1.07 for High RS diet compared to 1.00-1.04 for Intermediate RS diet) also remains to be discerned. As the High SDS diet had relatively low *x*50_*C*_ (carbohydrate oxidation mode, 0.98-1.02) compared to the other carbohydrate-dominant diets, this suggests that the RCS in this diet was more slowly delivered to the body, thus decreasing carbohydrate oxidation.

For *b*_*C*_, the High RS diet had a significantly higher value than the Sucrose and Intermediate SDS diet groups, but all other diets did not differ (Table 2). This finding suggests there was a narrower spread in RER values for the mode representing carbohydrate oxidation for the High RS diet than for the Sucrose and Intermediate SDS diets (Figure 3). As explained above, a narrowing in the range of RER values in either the carbohydrate or fat oxidation modes could be indicative of a swift and more complete change in substrate utilization. However, this parameter on its own does not indicate metabolic flexibility.

A main effect of the Mgam genotype was observed for the *x*50_*F*_, *b*_*F*_, *b*_*C*_, and *α* parameters (*p*=0.002, *p*=0.004, *p*<0.0001 and *p*=0.04, respectively), such that null mice had significantly higher *x*50_*F*_, *b*_*C*_, and *α* parameter estimates, but lower *b*_*F*_, than wild-type mice (Table 2). The overall higher *x*50_*F*_ (representing the fat oxidation mode) for null mice suggests that reduced ability to digest carbohydrate reduced fat oxidation, aligning with the concept that fat oxidation levels change more as a function of carbohydrate intake and energy expenditure than fat intake ^56^. Null mice burned less fat as fuel compared to wild-type mice. However, the higher *α* for null mice indicates more oxidation was occurring in the first mode (fat oxidation mode). Together, these results for *x*50_*F*_ and *α* suggest that null mice exhibited disrupted fat oxidation, specifically lingering at an incomplete level of fat oxidation (i.e., *x*50_*F*_ ∼0.81). This state may represent mixed metabolism of carbohydrate and fat, possibly indicative of metabolic inflexibility. The high *b*_*C*_ but low *b*_*F*_ for null mice show that their RER values were clustered more closely in the second mode, but more widely in the first mode. Overall, these differences were minor compared to what we anticipated for mice lacking a complete set of enzymes, but they support the previous findings that Si is able to handle the digestion of most types of starch-hydrolyzed products of α-amylase, such as those used here, while Mgam has wider versatility for hydrolyzing more difficult to digest starches and other α-glucans ^61^.

Regarding AMG supplementation, there was a main effect of cycle (i.e., with AMG vs. without AMG) for the *x*50_*F*_ and *x*50_*C*_, parameters (*p*<0.0001 and *p*=0.002, respectively; Table 2). For *x*50_*F*_, values were generally higher with AMG compared to without AMG, which implies that adding AMG shifted oxidation toward carbohydrate metabolism, leading to concurrent oxidation of carbohydrate and fat [aligning with the carbohydrate displacement theory of Flatt ^56^]. We reason that the departure from RER values representing ideal fat metabolism (RER=0.70) observed with AMG supplementation may be undesirable, given that a state of mixed metabolism has been proposed to reflect mitochondrial gridlock and has been associated with obesity-related perturbations in fuel usage ^6,62^. Additional factors to be considered when interpreting the implications of a state of mixed metabolism, when examined on a preprandial and postprandial scale, are diet composition and total amount consumed in a given time period. For *x*50_*C*_, adding AMG also increased *x*50_*C*_ values, signifying supplementation of the diets with AMG increased median RER of the second mode (representing carbohydrate oxidation). This suggests AMG increased partitioning of carbohydrates to be used for metabolism and energy use or storage. Taken together, these findings suggest that supplementation with AMG may provoke similar metabolic consequences to consumption of a readily available carbohydrate (e.g., sucrose, rapidly digestible starch) and, specifically, to the simultaneous oxidation of carbohydrate and fat that has been described as metabolic gridlock that is characteristic of metabolic inflexibility ^6^.

An interaction effect was found for *α* such that genotype × cycle was significant (*p*=0.03) and for *b*_*F*_ such that genotype × diet and genotype × cycle × diet (*p*=0.0009, *p*=0.009) were significant. Furthermore, for *x*50_*F*_ the interaction among genotype × cycle × diet was on the border of significance (*p*=0.05). No other interactions were significant (*p*>0.05).

### New approach to characterize metabolic flexibility

The protocol developed in the present work addresses a heretofore lack of a quantitative means to comprehensively assess metabolic flexibility using RER data. A unique aspect of PRCF is that it represents a cumulative percentage of the distribution. Therefore, when applying the Mixed Weibull Cumulative Distribution to the data and solving for a given *x* (RER value), the proportion of RER values in the dataset that fall below that unique *x* value will be the resulting *y* solution. Given this characteristic, an arithmetic approach can be used to calculate the proportion of the distribution falling between any two given *x* values. Accordingly, we propose that calculating the portion of RER values falling within the ideal ranges for carbohydrate oxidation and for lipid oxidation is an intuitive means to quantify metabolic flexibility, which we define as the Metabolic Flexibility Factor (MFF). This can be employed for different timescales, but for the present analyses we used a timescale of 24 h per condition. MFF represents the estimated proportion of values that fall within ±10% of the theoretical modes for fat and carbohydrate oxidation (RER = 0.7 and 1.0, respectively), and a higher MFF indicates greater metabolic flexibility (Eq. 3; Figure 2).

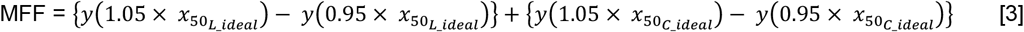

### Slow carbohydrate digestion enhances metabolic flexibility

Applying the MFF approach, there were main effects of diet, cycle (AMG), and the interaction between diet × cycle (*p*<0.0001, *p*=0.0002, and *p*=0.0008, respectively; Figure 5). *Post-hoc* tests revealed that the High SDS, Intermediate SDS, Intermediate RS, and High RS diets all had significantly higher MFF values than the Sucrose and High-fat diets. Further, Sucrose had the lowest MFF overall, significantly differing from the High-fat diet. The High-fat and Sucrose diets can be considered as ‘anchors’ among the diets tested – both had low MFF values and thus low metabolic flexibility, but in distinct ways. Namely, the High-fat diet led to a larger proportion of RER values falling between the modes of ideal carbohydrate and fat oxidation, whereas the Sucrose diet resulted in an overall shift toward greater carbohydrate oxidation (represented by higher RER for both modes) as well as RER values indicative of fat storage (RER > 1.0). In addition to this, and strikingly, the MFF for the High SDS diet was higher than that for High RS diet (*p*=0.04), which indicates that SDS consumed in sufficient quantity may be superior for metabolic health than RS. AMG supplementation, which increases starch digestion rate, significantly decreased MFF for the High RS diet (*p*=0.0006), whereas within all other diets it had no significant effect (*p*>0.05). These results collectively support the notion that diets high in SDS and RS result in superior metabolic flexibility compared to diets high in sucrose or fat. Furthermore, and importantly, the difference in MFF between the High SDS and High RS diets suggests that slow carbohydrate digestion enhances metabolic flexibility more than reduced/resistant carbohydrate digestion.

**Figure 5.**
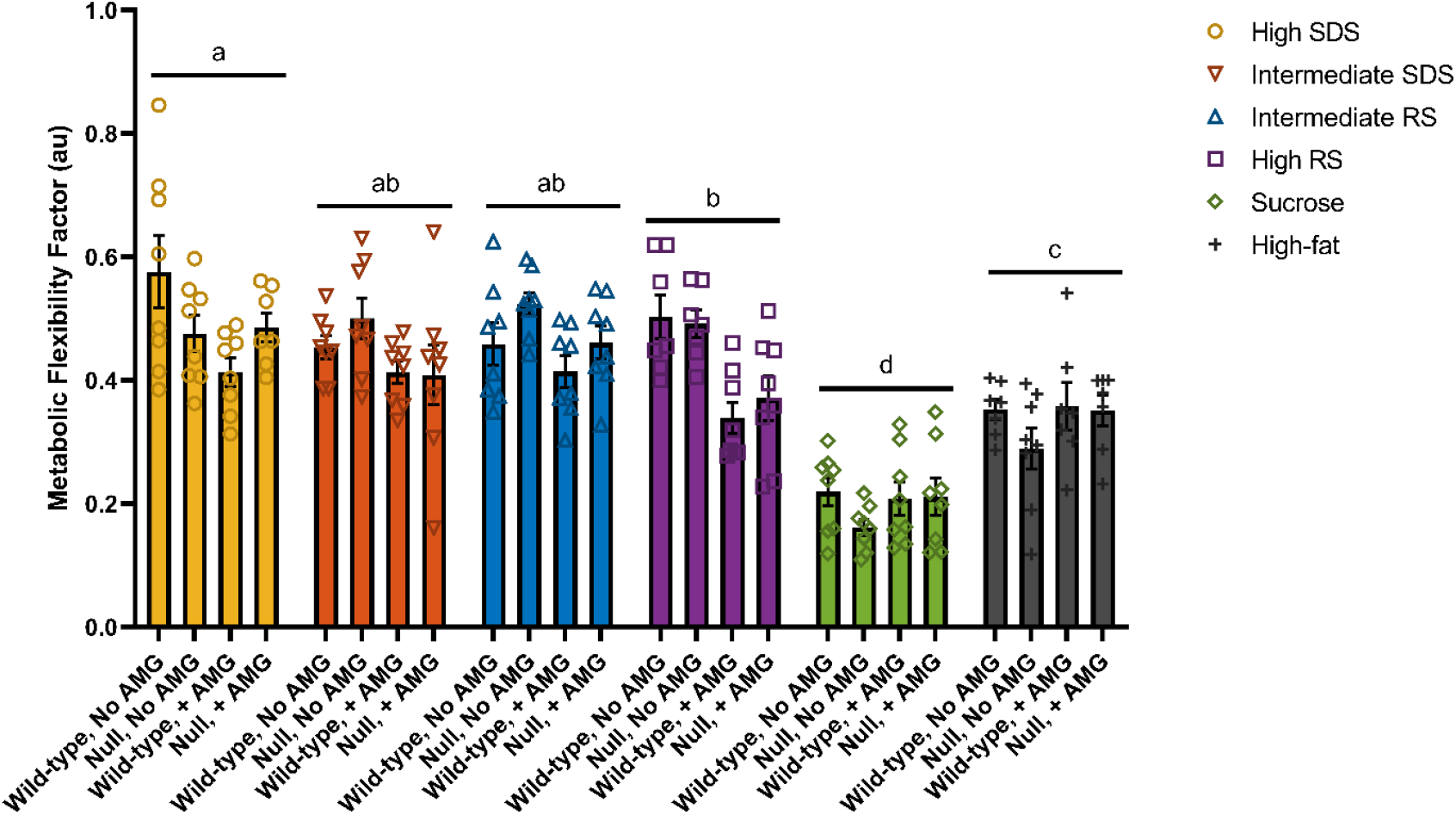
Mean Metabolic Flexibility Factor (MFF) for the diet × genotype × cycle [AMG] groupings. A high MFF is suggested to indicate improved metabolic flexibility. Bars represent mean values, shown with ± standard error of the mean, and symbols represent individual observations (mice) per dietary condition. Different letters indicate statistically significant differences (*p*<0.05) per diet. In addition to the main effects of diet shown (*p*<0.0001), there was also an effect of cycle [AMG], such that AMG decreased MFF compared to without AMG (*p*=0.0002). There was also an interaction effect for diet × cycle for MFF (*p*=0.0008). AMG, amyloglucosidase; au, arbitrary units; MFF, Metabolic Flexibility Factor.

## Discussion

The objective of this study was to determine how altering carbohydrate digestibility affects its utilization as a substrate for energy and the ensuing impact on metabolic flexibility between oxidation of carbohydrate and fat. We assessed six different diets and distinct dietary conditions as well as devised and carried out novel applications of mathematical modeling approaches to explore this objective.

For diet-related differences, diets containing a proportion of SDS (18%, 35%, or 53% levels of RCS), had superior ability to promote fat oxidation (lower *x*50_*F*_) and generally allowed for enhanced metabolic flexibility (MFF). Notably, the MFF for the High SDS diet was higher than for the High RS diet, suggesting that slow carbohydrate digestion increases metabolic flexibility, and that slow digestion provides greater impact than reduced or resistant digestion of starch. The High SDS diet did not promote carbohydrate oxidation (i.e., increase *x*50_*C*_). High *x*50_*C*_ values observed for the other carbohydrate-dominant diets may be detrimental because they exceeded 1.00 RER. In addition, the Sucrose diet had the highest carbohydrate oxidation (*x*50_*C*_) and the High-fat diet had the lowest carbohydrate oxidation (*x*50_*C*_), validating our modeling approaches. High metabolic inflexibility was indicated for the Sucrose diet and moderately indicated for the High-fat diet via the quantification of MFF.

Mice lacking intact Mgam function did not have lower carbohydrate oxidation (as would have been shown through low *x*50_*C*_ values; Figure 4, Table 2), but instead had disrupted fat oxidation. Si may have been upregulated in null mice ^34,36^, perhaps as compensation for the lack of maltase-glucoamylase. Despite these findings, the observed differences between null and wild-type mice were surprisingly minor overall, considering that the null mice lacked a complete set of starch digesting enzymes. Because the null mice with only Si generally performed as well as wild-type mice, this supports previous research that Si is a more dominant starch digesting enzyme than Mgam ^34,39,63^.

AMG, which increased starch digestion rate, generally increased carbohydrate oxidation, as indicated by higher *x*50_*C*_ values, for all but the Sucrose and High-fat diets. One potential concern with the AMG supplement is that it appeared to shift substrate utilization toward carbohydrate, but not representing fat oxidation (as indicated by the higher *x*50_*F*_ values and lack of differences in *b*_*C*_, and decreased MFF values), creating unfavorable conditions for complete switching between carbohydrate and fat substrates for oxidation.

Our study focused on oxidation, but implicit to this is a “toggling” between oxidation of absorbed glucose vs. fermentation of RS generating short-chain fatty acids. This topic requires further investigation.

Using novel approaches, the findings from this work indicate that moderated or slow carbohydrate digestion, such as observed for the High SDS diet, promotes fat oxidation and enables an improved ability to switch between complete carbohydrate oxidation and complete fat oxidation. This improvement in metabolic flexibility between utilization of substrates may have implications on deposition of adipose tissue, insulin sensitivity, and mitochondrial function in the body ^4,6^, even in the absence of differences in energy expenditure. These results indicate a new potential role of slowly digestible carbohydrates or moderated carbohydrate digestion rate in metabolic health, which could have practical application in the production of whole and processed foods to sustain a healthier, growing human population.

## Materials and Methods

All study procedures were conducted with full approval from the Baylor College of Medicine Institutional Animal Care and Use Committee (IACUC; protocol AN-1577).

### Animals

Equal groups of Mgam knockout (null) and wild-type mice were used (*n*=8 per group; 8 males in null group, 7 males and 2 females in wild-type group). Both groups were Sv/129 mice (The Jackson Laboratory, Bar Harbor, ME). For the null mice, the Mgam gene was ablated as reported previously ^36^ and genotyping proceeded using quantitative polymerase chain reaction (PCR) of tail DNA as described in Nichols et al. ^37^. For the wild-type mice, the Mgam gene was not ablated. All mice were weaned at least 42 days before initiation of experimentation. Mice were singly housed in rooms maintained at 22 ± 2ºC with a 12:12 h dark:light cycle (18:00-06:00 dark).

### Experimental diets

Mice were fed six different diets *ad libitum* over time in the following sequence: **1)** High SDS (53% Raw Corn Starch [RCS] diet; Envigo-Teklad TD.01629 containing raw corn starch; Envigo, Indianapolis, IN), **2)** High RS (53% High-Amylose Starch [HAS] HAS; Envigo-Teklad TD.02130 containing Novelose 260 high-amylose maize starch), **3)** Sucrose (65% Sucrose diet; Envigo-Teklad TD.02129), **4)** Intermediate RS (Envigo-Teklad TD.02130), **5)** Intermediate SDS (Envigo-Teklad TD.02130), **6)** High-fat (21.2% fat, or 42% kcal from fat; Envigo-Teklad TD.88137). Digestion profiles of the experimental diets are visualized in Figure 1. With the exception of the High-fat diet, the only difference among the diets was in the carbohydrate portion (Table S1). The % kcal and % by weight from protein, carbohydrate, and fat were matched for all but the High-fat diet. Further, all diets had an energy density of 3.6 kcal/g with the exception of the High-fat diet, which had 4.5 kcal/g. When the mice were not on an experimental diet, they were fed a chow lab diet (PicoLab Rodent Diet 20, 5053; 62% kcal from carbohydrate, 25% kcal from protein, 13% kcal from fat; LabDiet, St. Louis, MO). Total starch in each diet was determined using a total starch analysis kit (amyloglucosidase/alpha-amylase method; Total Starch Assay Kit [AA/AMG], K-TSTA-50A, Megazyme, Wicklow, Ireland). Amounts of rapidly digestible starch (RDS), slowly digestible starch (SDS), and resistant starch (RS) were determined using the Englyst assay ^44,45^ (Table S2), and percent amylose within the carbohydrate component of each diet was calculated (considering that normal corn starch contains 28% amylose and Novelose 260 contains 70% amylose; Table S2). Furthermore, food quotients (macronutrient oxidation ratio for food or, in this case, for different diets) were calculated for all of the diets assuming a quotient of oxidation of 1.0 for carbohydrates, 0.70 for fats, and 0.825 for proteins (Table S2). Discussion of these results can be found in the Supplementary Information file. According to IACUC requirements, diets were stored for no longer than 6 weeks before being provided to animals.

For experimentation, animals were initially subjected to the diets and indirect calorimetry chambers for 4 cycles (24 h per cycle) prior to data collection as a means of acclimation (Figure S2). Due to the nature of these experiments, one animal was housed per chamber/cage. Following these 4 cycles, data collection proceeded for 4 more cycles (24 h per cycle) in the indirect calorimetry chambers as described below. Animals were weighed before and after being housed in the chambers. At least 10 day/night cycles served as a washout period between diets. Mice were also given free access to drinking water at all times (either with or without AMG supplementation for data collection).

### Amyloglucosidase (AMG) supplementation

During the latter two cycles of the data collection period in the indirect calorimetry chambers (cycles 3 and 4), each animal’s drinking water was replaced by drinking water supplemented with 0.5% amyloglucosidase (AMG) from *Aspergillus niger* (2 mL AMG/400 mL drinking water; AMG 300 L, AMP30095, activity 300 U/mL; Novo Nordisk BioChem, North America, Inc., Franklinton, NC). This supplementation level was determined considering a standard AMG activity of 0.2 U/g digestible starch in mice, and by assuming animals would consume 2.5 g starch/d; taken together, these factors indicated a level of 0.5 U/d was desired, which was then incorporated with the activity level of the AMG per mL (3000 U/mL) to indicate 2 mL AMG per 400 mL drinking water was required (0.5%). Following the termination of cycle 4 for each diet data collection period, the drinking water was switched back to the unsupplemented version.

### Indirect calorimetry – Respiratory Exchange Ratio (RER)

The respiratory exchange ratio (RER) of each mouse for the 4-cycle data collection period was determined using an open-circuit indirect calorimeter (version 5.61, Oxymax, Columbus Instruments, Columbus, OH) designed to measure consumption of oxygen (V_O2_) and production of carbon dioxide (V_CO2_). This 4-cycle data collection period was initiated following the 4-cycle acclimation period (Figure S2). With 16 separate chambers, this system had the capacity to house 16 mice simultaneously. Calibration of the V_O2_ and V_CO2_ sensors as well as flow meters was performed immediately prior to the data collection period for each diet. Sampling of inlet air from each chamber ensued every 43 min during the data collection period, with a 30 s measure time and 120 s settle time (referencing method 2, interval 1). Using equations derived from mass balance across the chamber, V_O2_ and V_CO2_ were calculated. The Weir equation ^64^ was used to internally calculate energy expenditure for each mouse, with adjustments made for individual mouse body weight (before being placed in indirect calorimetry chamber for each diet) and RER. RER values were averaged per 24 h separately for cycles 2 and 3 per treatment group (factors: diet × genotype × cycle), as well as analyzed and modeled for pooled treatment groups overall or per individual mice.

### Calculation of Percent Relative Cumulative Frequency (PRCF) of RER

Untransformed RER data for individual mice for 24 h was used to calculate RER percent relative cumulative frequency (PRCF) per mouse genotype, diet, and cycle (cycle 2 [without AMG] vs. cycle 3 [with AMG]) according to the method of Riachi et al. ^55^. Briefly, this involved the following steps: (1) the RER data was sorted in ascending order, (2) an interval of increment was selected (i.e., 0.01), (3) the frequency of observations per interval within the range of all values was calculated, (4) the cumulative frequency was calculated, and (5) the cumulative frequency was expressed as a percentile curve. The analysis was performed for individual mice per diet, genotype, and cycle.

### Modeling of PRCF

Following calculation of PRCF, plots of RER (ascending order) vs. PRCF were fit to the Mixed Weibull Cumulative Distribution function (Eq. 1). This function represents a bimodal distribution.

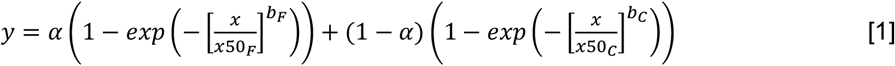

Where:

*y* = percent relative cumulative frequency (PRCF; 0 to 100%);

*α* = mixing weight parameter that represents the proportion of the first mode;

*x*50_*F*_ = median respiratory exchange ratio (median RER) for the <u>first</u> mode;

*b*_*F*_ = distribution breadth constant for <u>first</u> mode (dimensionless), indicative of slope for the first mode;

*x*50_*C*_ = median respiratory exchange ratio (median RER) for the <u>second</u> mode;

*b*_*C*_ = distribution breadth constant for the <u>second</u> mode (dimensionless), indicative of slope for the second mode.

To ensure that the first mode was associated with the lower values of RER, a constraint was imposed (Eq. 2, ^65^:

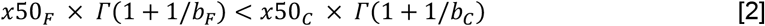

here *Γ* is the gamma function. Data were fit to Eq. 1 subject to the constraint in Eq. 2 using MATLAB R2022a with the function *fmincon*. The objective function was minimization of the sum of squared errors between the data and model.

Modeling was done using the “fitnlm” function with the nonlinear least squares method option in MATLAB (R2020a, Update 5, 9.8.0.1451342, The MathWorks, Inc., Natick, MA). The Mixed Weibull Cumulative Distribution function (Eq. 1) was used as in Drechsler and Ferrua ^65^. Bounds were placed on the *x*50_*F*_, and *x*50_*C*_ model parameters to restrict them to the range of RER values for each individual dataset, and *α* was restricted to the range of 0 to 1. Curve fits summarized per treatment group are shown in Figure S3, and all curve fits for individual animals for each dietary condition are shown in Figure S4-S9. Python and MATLAB codes to perform these curve fitting and data analyses are provided in the Supplementary Information file.

### Development and application of a new approach to quantify metabolic flexibility: Metabolic Flexibility Factor (MFF)

The Mixed Weibull Cumulative Distribution equation was evaluated, for each mouse per condition, for RER values within 10% of the ideal lipid and carbohydrate oxidation RER modes (0.7 ± 10% for lipid; 1.0 ± 10% for carbohydrate, Eq. 3).

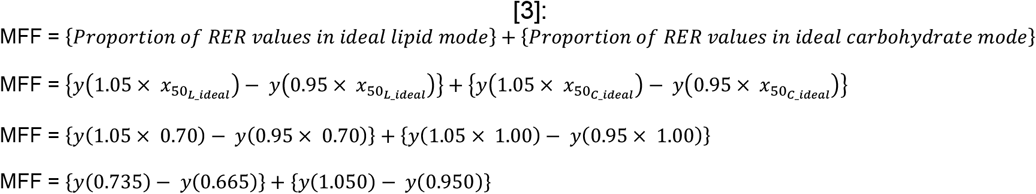

Where:

*y*(*x*) = Mixed Weibull Cumulative Distribution function solved for accompanying *x* (i.e., 0.665, 0.735, 0.950, 1.050), using the parameter estimates for *α, x*50_*F*_, *b*_*F*_, *x*50_*C*_, and *b*_*C*_ from curve fitting. MATLAB code to perform analysis can be found in the Supplementary Information file, and it can be employed using Dataset S1.

The resulting value from this equation represents the estimated proportion of values that fall within ±10% of the theoretical modes for fat and carbohydrate oxidation. To characterize true metabolic flexibility, a portion of values must fall within each mode. A higher MFF indicates greater metabolic flexibility.

### Mean RER per 24 h

Mean RER per 24 h cycle was calculated for each group (diet × genotype × cycle [without/with AMG]) by taking averages of the RER values for cycles 2 and 3.

### Body weight

The body weight of each mouse was measured prior to its placement into the indirect calorimetry chamber for each diet acclimation/experimental period for adjustment of the RER calculation (i.e., 6 times total, once before for each of the 6 diets). Additionally, body weight of each mouse was measured at the termination of each diet experimental period in the indirect calorimetry chamber system (i.e., after cycle 4). Results are reported in Supplementary Information.

### Statistical analysis

To determine the appropriate sample size of mice to use (diet × genotype × cycle; 24 treatment groups total [with 6 diets, 2 genotypes, and 2 cycles]), a power calculation was conducted based on average RER over a 24 h period (G*Power v.3.1.9.7) ^66^. The effect size was set at 0.34 (effect size *F*, ratio of population standard deviations) using the G*Power built-in determination tool by inputting the results of Fernández-Calleja et al. ^14^ for RER over a 24 h period. For six diets split into 24 treatment groups and power of 0.8, it was determined that a minimum of 5 mice was required per treatment group (*n*=5). Eight mice were used per treatment group (*n*=8).

SAS version 9.4 (SAS Institute, Cary, NC) was used to conduct all statistical analyses. Three-way ANOVA (PROC MIXED) with diet, mouse genotype, and cycle (with or without AMG) as fixed effects was used to determine differences in Mixed Weibull Cumulative Distribution function parameters (i.e., *α, x*50_*F*_, *b*_*F*_, *x*50_*C*_, and *b*_*C*_) as well as Metabolic Flexibility Factor. Residuals of all models were plotted and assessed for homoscedasticity and normality using the Brown-Forsythe test and Kolmogorov-Smirnoff test, respectively. Data were transformed as needed until they passed these tests for homoscedasticity and normally. Statistically significant differences were considered at *p*<0.05, and Tukey’s *post hoc* test for multiple comparisons was performed when the overall model was significant for main effects or interactions (*p*<0.05 for *F* value).

## Notes

### Competing Interest Statement

The authors have declared no competing interest.

