## Supplementary material for "Moderating carbohydrate digestion rate promotes metabolic flexibility in mice": Figure S1, Figure S2, Figure S3, Figure S4, Figure S5, Figure S6, Figure S7, Figure S8, Figure S9, Table S1, Table S2

**This PDF file includes:**

Supplementary Information Text (including Python code and MATLAB code)
Figures S1 to S9
Tables S1 to S2
Supplementary Information References

#### Supplementary Information Text

##### *Starch digestibility, percent amylose, and food quotient of experimental diets*

The general compositions of the six experimental diets are shown in Table S1. According to the results from the Englyst assay, the high slowly digestible starch (SDS) diet had the highest amount of rapidly digestible starch (RDS) and a negligible amount of resistant starch (RS) relative to the other diets (Table S2). The incrementally increasing amounts of RS were found according to the level of high-amylose starch (HAS, Novelose 260) inclusion in the remaining starch-based diets (Intermediate SDS, Intermediate RS, and High RS diets had 10.2, 15.3, and 28.5% RS, respectively; Table S2). It is important to note that Novelose 260 is not entirely comprised by RS, which accounts for the discrepancies between the Novelose 260 inclusion levels and actual amounts of RS in these diets. Additionally, Novelose 260 contains a fraction of pre-treated/pre-gelatinized starch that is likely very highly digestible. Although technically the diets containing Novelose 260 had lower percentages of rapidly digestible starch (RDS) compared to the High SDS diet, the RDS they did contain may have been highly digestible *in vivo* – in such a manner that is not accurately reflected *in vitro*. This speaks to one of the limitations of the Englyst assay (1, 2). The Sucrose, High-fat, and PicoLab (non-experimental) diets contained negligible amounts of RS. The PicoLab diet contained the highest amount of SDS among all the diets, which may be attributed to the inclusion of ground raw corn in this diet instead of corn starch (as was used for the experimental diets). Additionally, the corn starch used in the experimental diets was in the raw state. Raw corn starch (RCS) has a well-characterized slow digestion property (3), and, accordingly, the High SDS diet had a relatively high amount of SDS according to the Englyst assay.

The 42% fat (High-fat) diet was designed to mimic the average U.S. adult human diet in terms of macronutrient percentage and general composition. Its inclusion in these experiments not only served as a means to “calibrate” our results but also to add context for other studies using this diet.

A portion of maltodextrin was included in each of the diets because it improved their pelletization. This contributed to the fairly large fraction of rapidly digestible starch (55.1% to 74.0%, dry starch basis; Table S2) in the diets. Additionally, the High-fat diet contained more sucrose than starch (341.5 g/kg vs. 150 g/kg) and no maltodextrin; sucrose improves pelletization of the diet as well and thus no maltodextrin was required in the High-fat diet.

When percent amylose within the digestible carbohydrate component of each diet was calculated, the diets fell in the following order based on highest to lowest amylose percentage: High RS diet > Intermediate RS diet > Intermediate SDS diet > High SDS > PicoLab diet (non-experimental) > High-fat diet > Sucrose diet (Table S2). The spread in amylose percentages ranged from 0 to 57% of the carbohydrate component. It is worth noting that not all amylose constitutes RS (4), as there are numerous other chemical, structural, and physical factors that affect RS content.

Food quotients were calculated based on the percent of each macronutrient by weight in each of the diets and were very similar among the diets, ranging from 0.73 to 0.79 (Table S2). These calculations are solely based on macronutrient compositions (i.e., percentages of carbohydrate, protein, and fat). Such quotients may benefit from factoring in additional aspects of the diets, such as carbohydrate digestibility.

##### *Body weight*

Mice maintained stable body weight during their periods in the indirect calorimetry chambers for each diet ($p<0.05$; data not shown), which indicates that weight gain or loss did not contribute to differences in RER. There are a number of previous studies and reviews that take thorough approaches to modeling metabolism and body weight dynamics in mice and humans (5, 6). For the current study, we decided to reduce the scope to specifically examine RER as an indication of substrate utilization in the body, especially given that body weight did not change during the experimental periods.

***Strengths and potential limitations***

Novel approaches were used to examine how alterations in carbohydrate digestion affect carbohydrate oxidation in mice. Diets incorporating carbohydrate components with varying digestibilities were used, mice lacking one of the α-glucosidases that hydrolyzes starch degradation products were used (compared to wild-type mice that possessed the complete set of starch digestion enzymes) to further slow digestibility, and AMG was used to augment starch digestion. Including an acclimation period (four 24-h cycles) to the diet and indirect calorimetry chamber for each diet treatment helped ensure any differences observed were not due to initial exposure to the experimental conditions imposed. Integrating previously proposed approaches for analyzing RER with new approaches provided an innovative means to study dynamics in RER to examine the effects of different conditions on metabolism.

As for limitations, one notable constraint is that food intake was not measured. However, previous studies found that mice food intake in home cages vs. indirect calorimetry chambers did not differ (7), and in a previous experiment the food intake between the same type of null mice in this study (Mgam knockout) and wild-type mice did not differ for a carbohydrate-predominant diet (data not shown). It is possible that mice consumed more food for certain diets (e.g. Sucrose diet, High-fat diet) than others (e.g. High RS diet) because of diet palatability or appetite. We also did not measure activity level of the mice while they were in the indirect calorimetry chambers (the system used was not equipped to do so). We believe differences in activity level for individual mice may have contributed to some of the variation observed within treatment groups. Two assumptions related to interpretation of these experimental findings were that there were no changes in mitochondrial number or function for the different treatments and that the mouse is an appropriate model for human starch digestion. Another limitation in this work is that we did not include a treatment arm of gelatinized starch or another form or rapidly digestible starch (RDS), especially considering that humans regularly consume gelatinized starch as opposed to raw starch. As the diets tested contained substantial amounts of SDS and RS, incorporating a diet containing a greater proportion of RDS (e.g., maltodextrin) would have strengthened the study.

***Resource to replicate/employ the proposed mathematical modeling and analysis – Codes are provided here to conduct the same analysis using either MATLAB or Python***

MATLAB code for modeling using the Mixed Weibull Cumulative Distribution function to fit percent relative cumulative frequency (PRCF) data for RER and to calculate Metabolic Flexibility Factor (MFF) is shown in blue below. An example RER dataset is provided within the code.

### calculate_mff_python

#### *Steps:*

1. Start with Respiratory Exchange Ratio data (RER) and then calculate Percent Relative Cumulative Frequency (PRCF)
2. Conduct a curve fit of PRCF values using the Mixed Weibull function
3. Calculate the Metabolic Flexibility Factor (MFF)
4. Make a plot showing the data points and the model

In [71]:

```
# Get libraries
import numpy as np
import scipy as sp
import matplotlib.pyplot as plt
import scipy.optimize as opt
from scipy.special import gamma
```

#### *Step 1: Start with RER and calculate PRCF*

In [72]:

```
# Get RER data
RER = np.array([0.7, 0.71, 0.71, 0.72, 0.72, 0.72, 0.74, 0.69, 0.7, 0.72,
0.72, 0.71,
    0.78, 0.75, 0.97, 1.01, 0.95, 1.02, 0.99, 1, 1.05, 1.04, 0.97, 0.99,
    0.98, 1, 0.96, 1.03, 0.99, 1, 0.99, 0.88, 0.95, 0.92])

# Sort RER values from low to high
RER = sorted(RER)

# Next, convert RER data to an integer by multiplying by 100
RER = np.multiply(RER, 100)
RER = RER.astype('int')

# Use an increment of 1, which is 0.01 in the original scale
RERmesh = np.arange(np.amin(RER), np.amax(RER)+1, 1)

# For each increment, find the corresponding number of RER values
NValsInRange = []
NvalsTemp = []
for i in range(0, len(RERmesh)):
    NvalsTemp = len(RER[RER == RERmesh[i]])
    NValsInRange.append(NvalsTemp)

# Next find the cumulative number of values in each range
cumNValsInRange = np.cumsum(NValsInRange)
```

```
# Rescale RER to roughly 0-10
# Using the defaults in this code we get good curve fits when RER is roughly
0-10
# Divide by 10 later to go back to the original scale (roughly 0-1)
RERmesh = RERmesh/10

# Now calculate PRCF
# PRCF(x) is the cumulative percentage of all values below x
PRCF = np.divide(cumNValsInRange, cumNValsInRange[-1])

print(PRCF)
[0.02941176 0.08823529 0.17647059 0.32352941 0.32352941 0.35294118
 0.38235294 0.38235294 0.38235294 0.41176471 0.41176471 0.41176471
 0.41176471 0.41176471 0.41176471 0.41176471 0.41176471 0.41176471
 0.41176471 0.44117647 0.44117647 0.44117647 0.44117647 0.47058824
 0.47058824 0.47058824 0.52941176 0.55882353 0.61764706 0.64705882
 0.76470588 0.85294118 0.88235294 0.91176471 0.94117647 0.97058824
 1.        ]
```

#### ***Step 2: Curve fit using the mixed weibull function***

In [73]:

```
x = RERmesh
y = PRCF

# Define the Mixed Weibull function
def mixedWeibull(x, a, x50f, bf, x50c, bc):
    return a*(1-np.exp(-(x/x50f)**bf)) + (1-a)*(1-np.exp(-(x/x50c)**bc))

# Set initial guess
ig = np.array([0.25, 7, 15, 9, 20]) # [a, x50f, bf, x50c, bc]

# Define objective function
def SSE(beta, y, x):
    a, x50f, bf, x50c, bc = beta
    return np.sum((y - mixedWeibull(x, a, x50f, bf, x50c, bc))**2)

# Define the gamma constraint
def gammaConstraint(beta):
    a, x50f, bf, x50c, bc = beta
    return beta[3]*gamma(1+1/beta[4]) - beta[1]*gamma(1+1/beta[2])

# Set parameter bounds
bnds = ((0,1), (0,1e3), (0,1e3), (0,1e3), (0,1e3))

# Set the nonlinear constraint
con = {'type': 'ineq', 'fun': gammaConstraint}

# Conduct the curve fit
r = opt.minimize(SSE, ig, args=(y, x), bounds = bnds, constraints=con)

# Select the individual parameters
a = r.x[0]
x50f = r.x[1]
bf = r.x[2]
```

```
x50c = r.x[3]
bc = r.x[4]

# x50f and x50c should be divided by 10 to return the original scale
betaHat = [a, x50f/10, bf, x50c/10, bc]

print(betaHat)

# Calculate R squared for the fit
yHat = mixedWeibull(x, a, x50f, bf, x50c, bc)

def rSquared(y, yHat):
# This function calculates R squared from data (y) and model values (yHat)
    resid = y - yHat
    SSresid = np.sum(resid**2)
    SStotal = np.sum((y-np.mean(y))**2)
    return 1-(SSresid/SStotal) #rsquared

rsq = str(round(rSquared(y,yHat),3))
print("R squared = ", rsq)
```

```
[0.40079615297496374, 0.7174086536756022, 51.43948307756947,
0.9952501567637277, 30.235927817053255]
R squared =  0.992
```

### Step 3: Calculate MFF

In [74]:

```
# MFF defines how much of the increase in PRCF occurs within 10% of the
# ideal RER value of fat digestion (0.7) OR within 10% of the idea RER
# value of carbohydrate digestion (1.0)

# Find the upper and lower bounds of the MFF region
# For carbs, this comes to 0.665-0.735
# For fat, this comes to 0.95-1.05

# Recall the x values for the curve fit were multiplied by 10
# So these limits also need to be multiplied by 10

# Calculate y values at the boundaries of the carb digestion region
yCarbLower = mixedWeibull(0.665*10, a, x50f, bf, x50c, bc)
yCarbUpper = mixedWeibull(0.735*10, a, x50f, bf, x50c, bc)

# Calculate y values at the boundaries of the fat digestion region
yFatLower = mixedWeibull(0.95*10, a, x50f, bf, x50c, bc)
yFatUpper = mixedWeibull(1.05*10, a, x50f, bf, x50c, bc)

# MFF is simply the sum of the increase that occurs in each region
MFF = yCarbUpper-yCarbLower + yFatUpper-yFatLower
print(MFF)
```

```
0.8456604672751341
```

### Step 4: Plot the data and model

In [75]:

```
# Plot the data
#plt.scatter(x/10, PRCF, color = 'black', label='Data')
```

```
plt.scatter(x/10, PRCF, facecolors='none', edgecolors='black', label='Data')

# Axis labels
plt.ylabel('Percent Relative Cumulative Frequency (PRCF, %)')
plt.xlabel('Respiratory Exchange Ratio (RER)')

# Set axis limits
plt.ylim([0, 1.1])
plt.xlim([0.6, 1.1])

# Evaluate the function on a fine mesh for plotting
xModel = np.arange(6, 11, 1e-3)
yModel = mixedWeibull(xModel, a, x50f, bf, x50c, bc)

# Plot the model
plt.plot(xModel/10, yModel, color = 'red', lw = 2, label='Model fit, R^2 =
'+rsq)

# Add a legend
plt.legend()

# Put MFF in  the title
plt.title("MFF = " + str(round(MFF,3)))

plt.show()
```

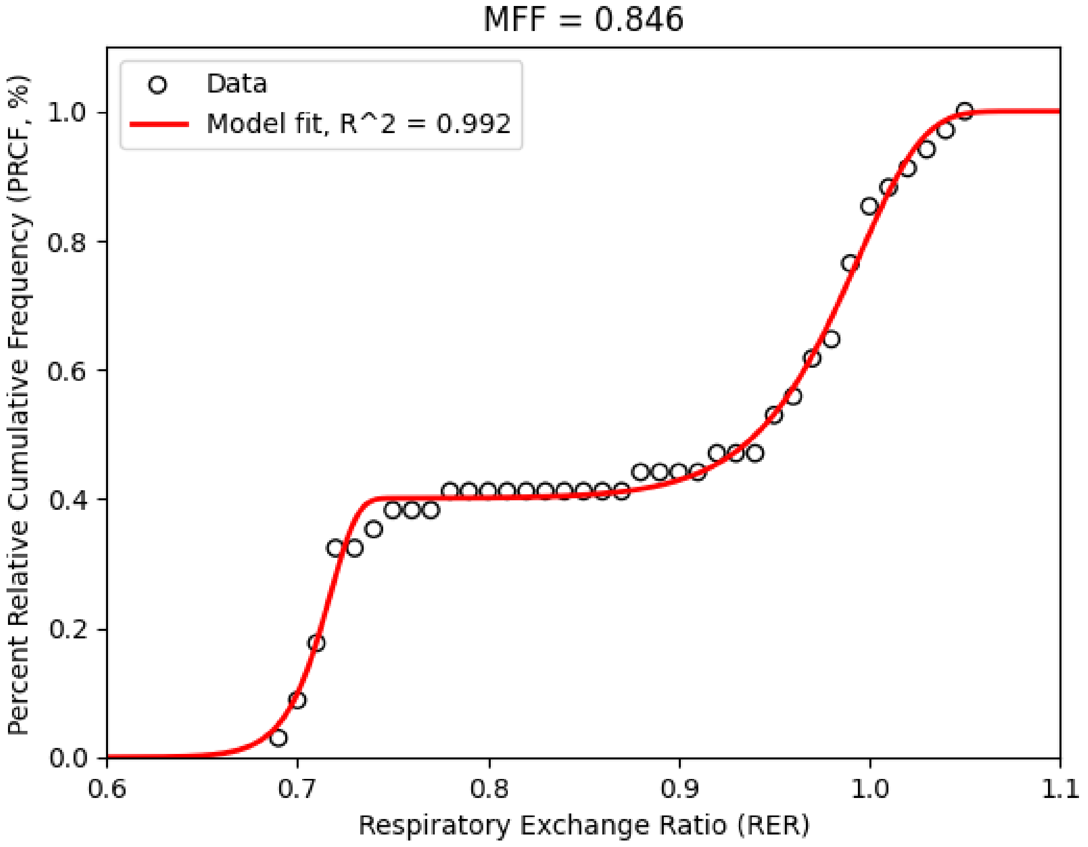

### calculate_mff_matlab

#### *Steps:*

1. Start with Respiratory Exchange Ratio data (RER) and then calculate Percent Relative Cumulative Frequency (PRCF)
2. Conduct a curve fit of PRCF values using the Mixed Weibull function
3. Calculate the Metabolic Flexibility Factor (MFF)
4. Make a plot showing the data points and the model

### Step 1: Start with RER and calculate PRCF

```
% Get RER data
RER = [0.7; 0.71; 0.71; 0.72; 0.72; 0.72; 0.74; 0.69; 0.7; 0.72; 0.72; 0.71;
...
    0.78; 0.75; 0.97; 1.01; 0.95; 1.02; 0.99; 1; 1.05; 1.04; 0.97; 0.99;...
    0.98; 1; 0.96; 1.03; 0.99; 1; 0.99; 0.88; 0.95; 0.92];

% Sort RER values from low to high
RER = sort(RER);

% Next, convert RER data to an integer by multiplying by 100
RER = RER*100;

% Use an increment of 1, which is 0.01 in the original scale
RERmesh = min(RER):1:max(RER);

% For each increment, find the corresponding number of RER values
for i = 1:length(RERmesh)
    NValsInRange(i) = length(RER(RER==RERmesh(i)));
end

% Next find the cumulative number of values in each range
cumNValsInRange = cumsum(NValsInRange);

% Rescale RER to roughly 0-10
% Using the initial guess in this code we get good curve fits when RER is
roughly 0-10
% Divide by 10 later to go back to the original scale (roughly 0-1)
RERmesh = RERmesh/10;

% Now calculate PRCF
% PRCF(x) is the cumulative percentage of all values below x
PRCF = cumNValsInRange./cumNValsInRange(end)
```

### Step 2: Curve fit using the mixed weibull function

```
x = RERmesh;
y = PRCF;

% Set initial guess
ig = [0.25, 7, 15, 9, 20]; % [a, x50f, bf, x50c, bc]
```

```
% Define objective function (minimize sum of squared errors, SSE)
objFun = @(beta) sseMixedWeibull(beta, x, y);

% Leave inequality constraints empty
Aineq = [];
bineq = [];

% Leave equality constraints empty
Aeq = [];
beq = [];

% Set parameter bounds
lb = [0, 0, 0, 0, 0]; % lower bounds = [a, x50f, bf, x50c, bc]
ub = [1,1e3,1e3,1e3,1e3]; % upper bounds = [a, x50f, bf, x50c, bc]

% Set nonlinear constraint
nonlcon = @gammaConstraint;

% Set options
options = optimoptions('fmincon','display','none');

% Conduct the curve fit
betaHat = fmincon(objFun, ig, Aineq, bineq, Aeq, beq, lb, ub, nonlcon,
options);

% Select the individual parameters from the output
a = betaHat(1);
x50f = betaHat(2);
bf = betaHat(3);
x50c = betaHat(4);
bc = betaHat(5);

% x50f and x50c should be divided by 10 to return the original scale
betaHat = [betaHat(1), betaHat(2)/10, betaHat(3), betaHat(4)/10, betaHat(5)]
```

### Step 3: Calculate MFF

```
% MFF defines how much of the increase in PRCF occurs within 10% of the
% ideal RER value of fat digestion (0.7) OR within 10% of the idea RER
% value of carbohydrate digestion (1.0)

% Find the upper and lower bounds of the MFF region
% For carbs, this comes to 0.665-0.735
% For fat, this comes to 0.95-1.05

% Recall the x values for the curve fit were multiplied by 10
% So these limits also need to be multiplied by 10

% Calculate y values at the boundaries of the carb digestion region
yCarbLower = mixedWeibullFunction(a, 0.665*10, x50f, bf, x50c, bc);
yCarbUpper = mixedWeibullFunction(a, 0.735*10, x50f, bf, x50c, bc);

% Calculate y values at the boundaries of the fat digestion region
yFatLower = mixedWeibullFunction(a, 0.95*10, x50f, bf, x50c, bc);
yFatUpper = mixedWeibullFunction(a, 1.05*10, x50f, bf, x50c, bc);
```

```
% MFF is simply the sum of the increase that occurs in each region
MFF = yCarbUpper-yCarbLower + yFatUpper-yFatLower
```

#### Step 4: Plot the data and model

```
% Plot the data
scatter(RERmesh/10, PRCF, 'ko', 'LineWidth', 1.5);

% Axis labels
ylabel('Percent Relative Cumulative Frequency (PRCF, %)')
xlabel('Respiratory Exchange Ratio (RER)')

% Set axis limits
ylim([0, 1.1])
xlim([0.6, 1.1])

hold on
box on

% Get yHat: predicted y values
yHat = mixedWeibullFunction(a, x, x50f, bf, x50c, bc);

% Calculate r squared
rsq = rSquared(y, yHat);

% Evaluate model on a finer mesh for a smoother plot
xModel = linspace(6, 11, 1e3);
yModel = mixedWeibullFunction(a, xModel, x50f, bf, x50c, bc);

% Plot the model
pModel = plot(xModel/10, yModel, '-', 'LineWidth', 1.5, 'Color', 'r');

% Rsquared and MFF values converted to text
rsqTxt = num2str(round(rsq,3));
MFFTxt = num2str(round(MFF,3));

% Add a legend
leg = legend('Data', strcat('Model fit: R^2 = ','{ }', rsqTxt));
leg.Location = 'NorthWest';

% Put MFF in the title
title(strcat('MFF = ', '{ }', MFFTxt))
```

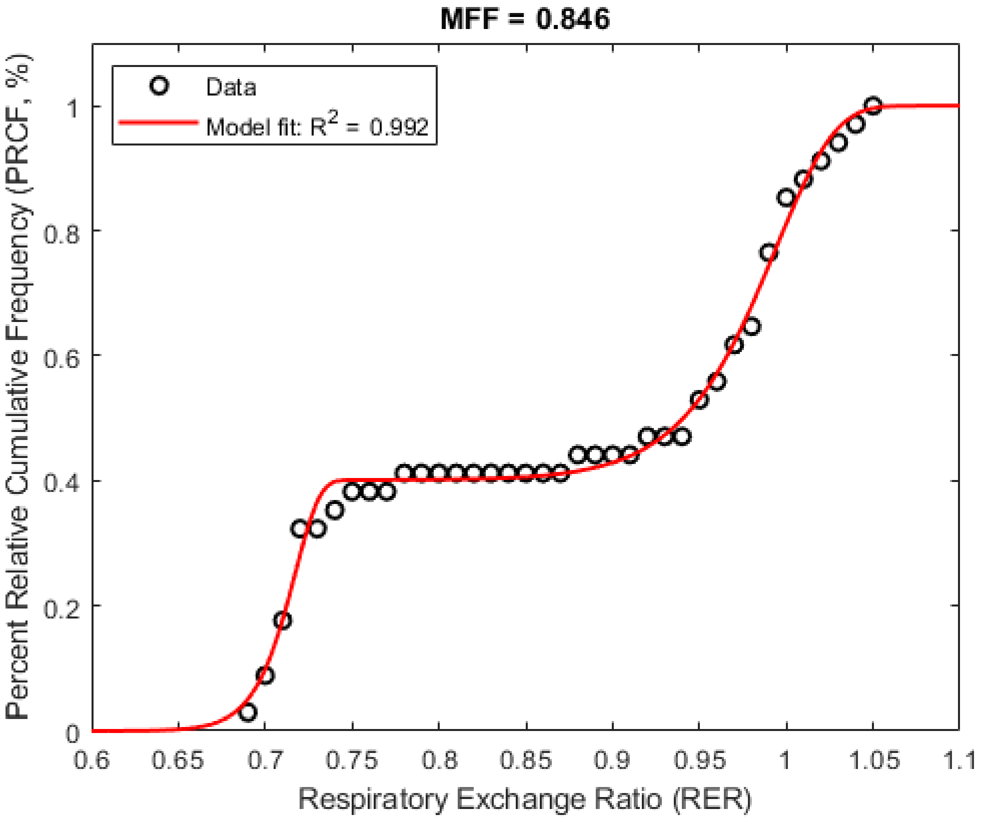


### Functions

```
FUNCTION: Define the mixed weibull function
function yHat = mixedWeibullFunction(a, x, x50f, bf, x50c, bc)
% This is the function whose parameters we will change to try to find a good
curve fit to the data

    yHat =  a*(1-exp(-(x/x50f).^bf)) + (1-a)*(1-exp(-(x/x50c).^bc));
end

FUNCTION: objective function for the curve fit
function sse = sseMixedWeibull(betaHat, x, y)
% This is the objective function for the mixed weibull fit

% Minimizing this function gives betaHat, the best parameters
% for the mixed weibull function when fitting to yData

% The criterion is sum of squared errors (SSE)

    % betaHat = [a, x501, b1, x502, b2]
    a = betaHat(1);
    x50f = betaHat(2);
    bf = betaHat(3);
    x50c = betaHat(4);
    bc = betaHat(5);

    yHat = mixedWeibullFunction(a, x, x50f, bf, x50c, bc);
```

```
    % Calculate sum of squared errors
    sse = sum(norm(yHat-y).^2);
end

FUNCTION: calculate rsquared
function rsq = rSquared(y, yHat)
% This function calculates R squared from data (y) and model values (yHat)

    resid = y - yHat;
    SSresid = sum(resid.^2);
    SStotal = sum((y-mean(y)).^2);
    rsq = 1-(SSresid/SStotal); % rsquared
end

FUNCTION: define a nonlinear constraint for the curve fit
function [c, ceq] = gammaConstraint(betaHat)
% This function imposes a nonlinear constraint on the curve fit

% For more information please see the following reference:
% Drechsler and Ferrua, 2016, "Modelling the breakdown mechanics of solid
% foods during gastric digestion", Food Res. Int,
10.1016/j.foodres.2016.02.019

% For more information on nonlinear constraints in MATLAB:
% https://www.mathworks.com/help/optim/ug/nonlinear-constraints.html

    % betaHat = [a, x501, b1, x502, b2]
    x50f = betaHat(2);
    bf = betaHat(3);
    x50c = betaHat(4);
    bc = betaHat(5);

    c = x50f*gamma(1+1/bf) - x50c*gamma(1+1/bc);
    ceq = [];
end
```

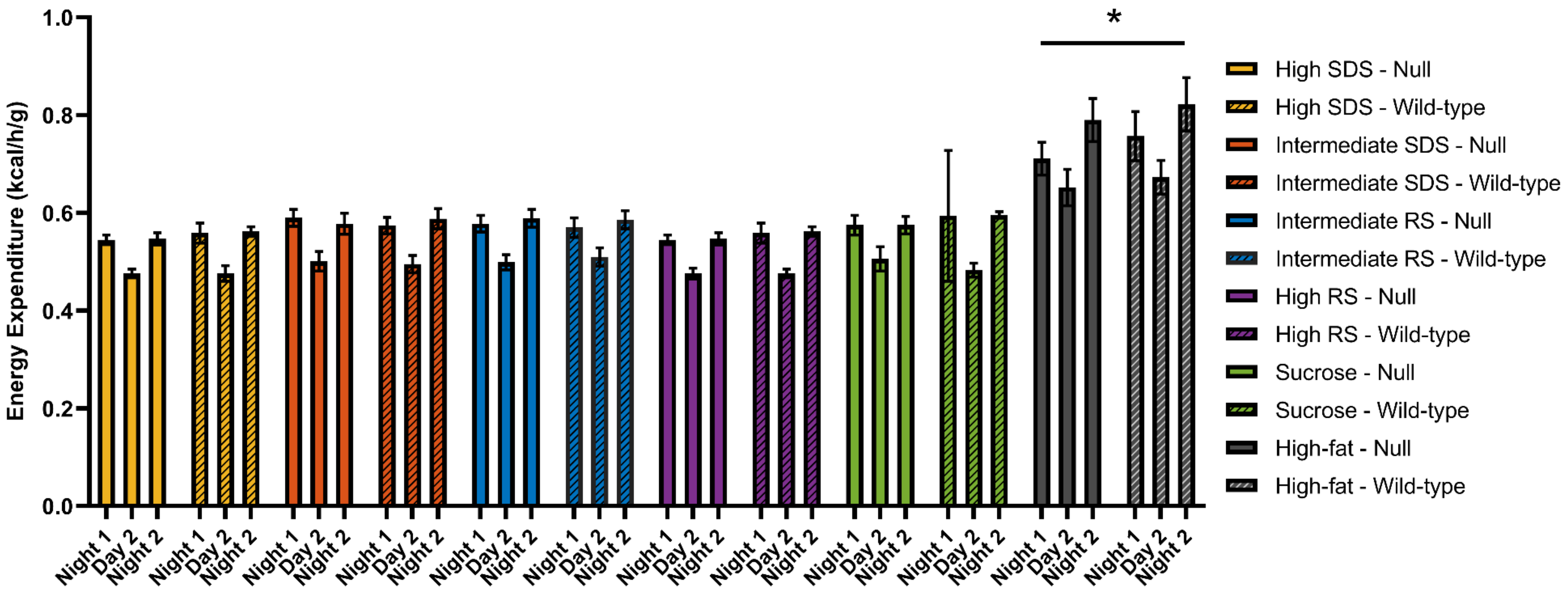


**Fig. S1.** Energy expenditure split by 12-h periods (day/night) for null and wild-type mice fed 6 different diets (*n*=8 each group). Error bars represent ± standard error of the mean (SEM). Null indicates maltase-glucoamylase (Mgam) knockout mice.

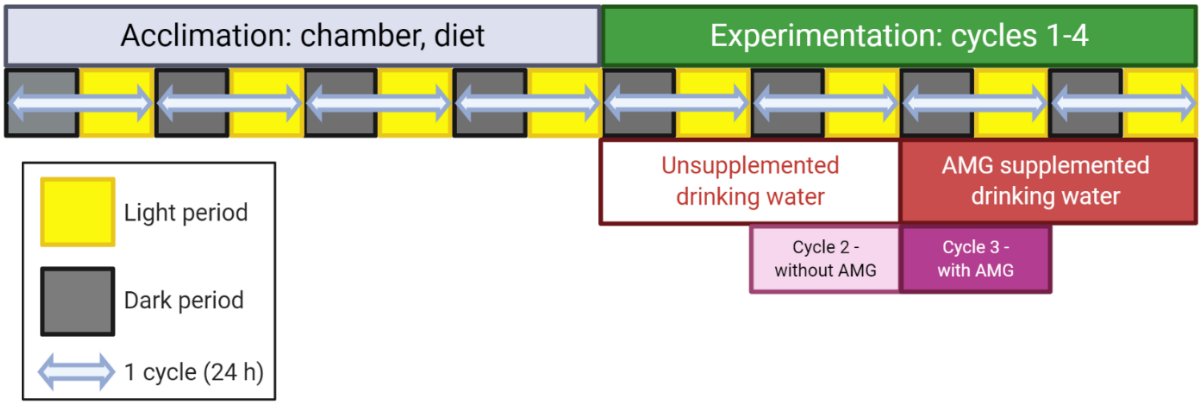


**Figure S2.** Diagram of indirect calorimetry experiments protocol per diet. Figure made using BioRender. AMG, amyloglucosidase.

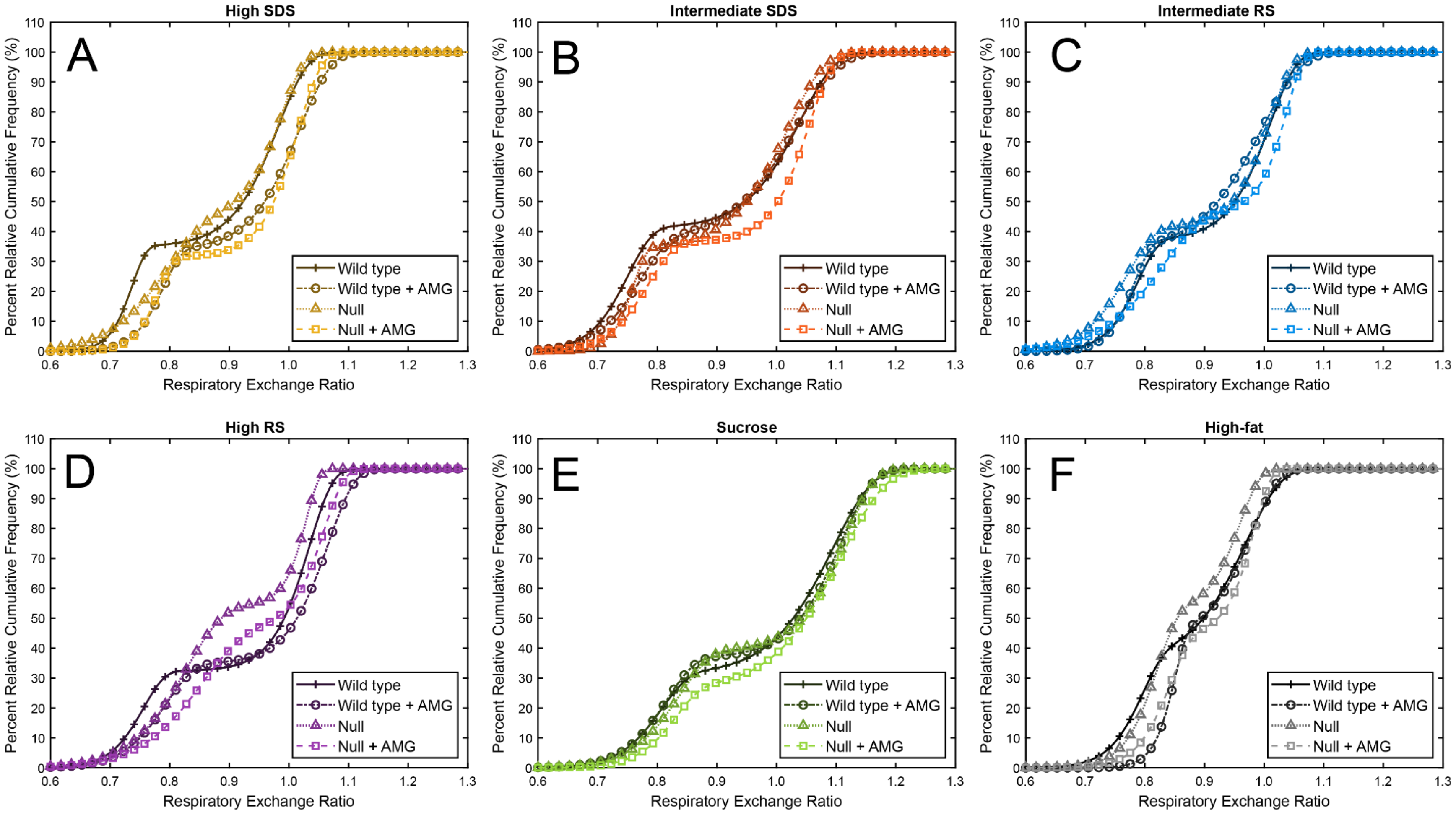


**Fig. S3.** Percent relative cumulative frequency (PRCF) curves for RER data from individual mice per diet (diet × mouse genotype × cycle). Split per diet: High SDS (A), Intermediate SDS (B), Intermediate RS (C), High RS (D), Sucrose (E), High-fat (F). Null indicates maltase-glucoamylase (Mgam) knockout mice. AMG, amyloglucosidase.

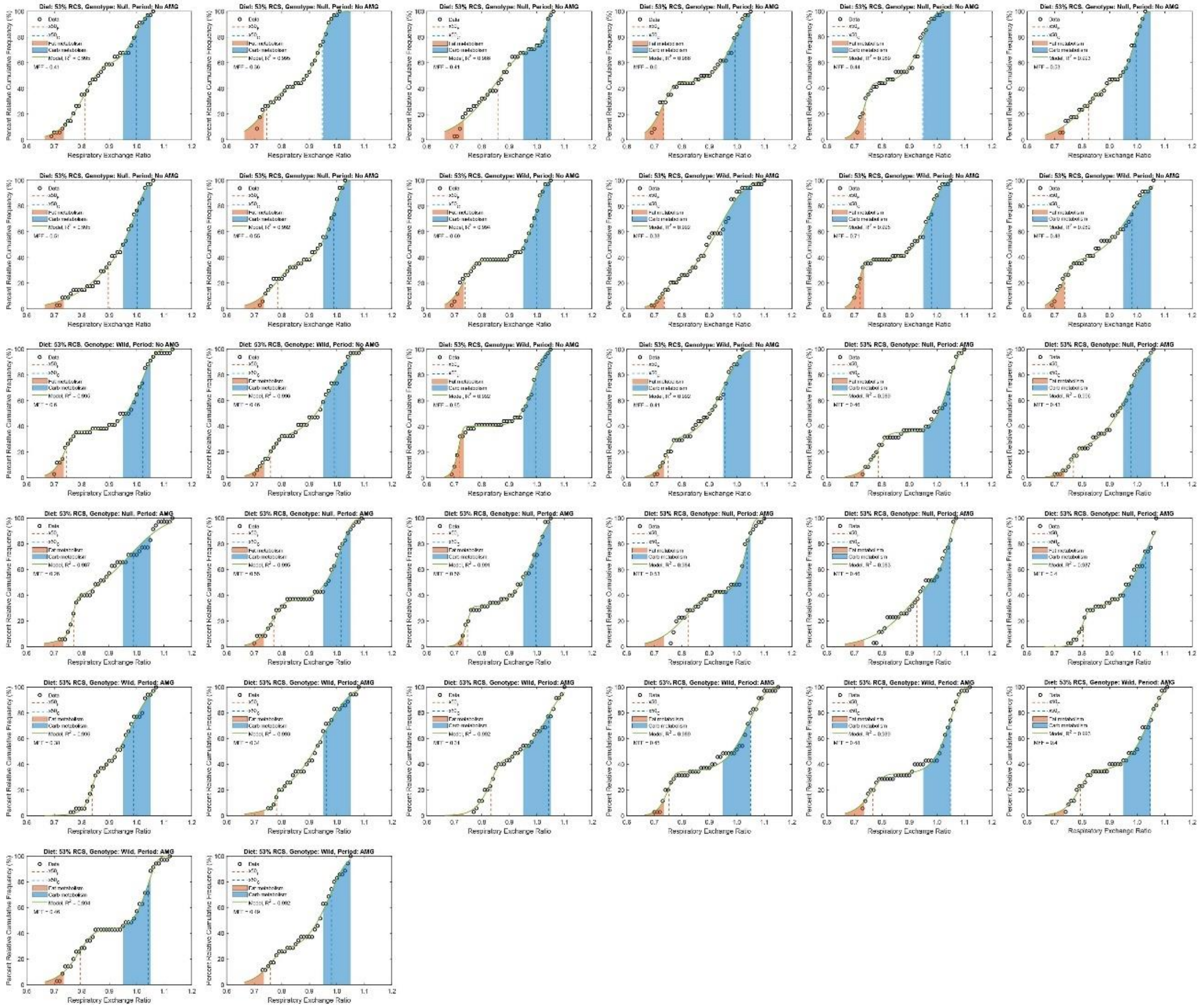

**Fig. S4.** Data and curve fits for Percent Relative Cumulative Frequency (PRCF) modeling using the Mixed Weibull Cumulative Distribution function for all individual mice on the High SDS diet. Red shading represents the ‘ideal’ fat oxidation range of respiratory exchange ratio (RER), while the blue shading represents the ‘ideal’ carbohydrate range of RER values.

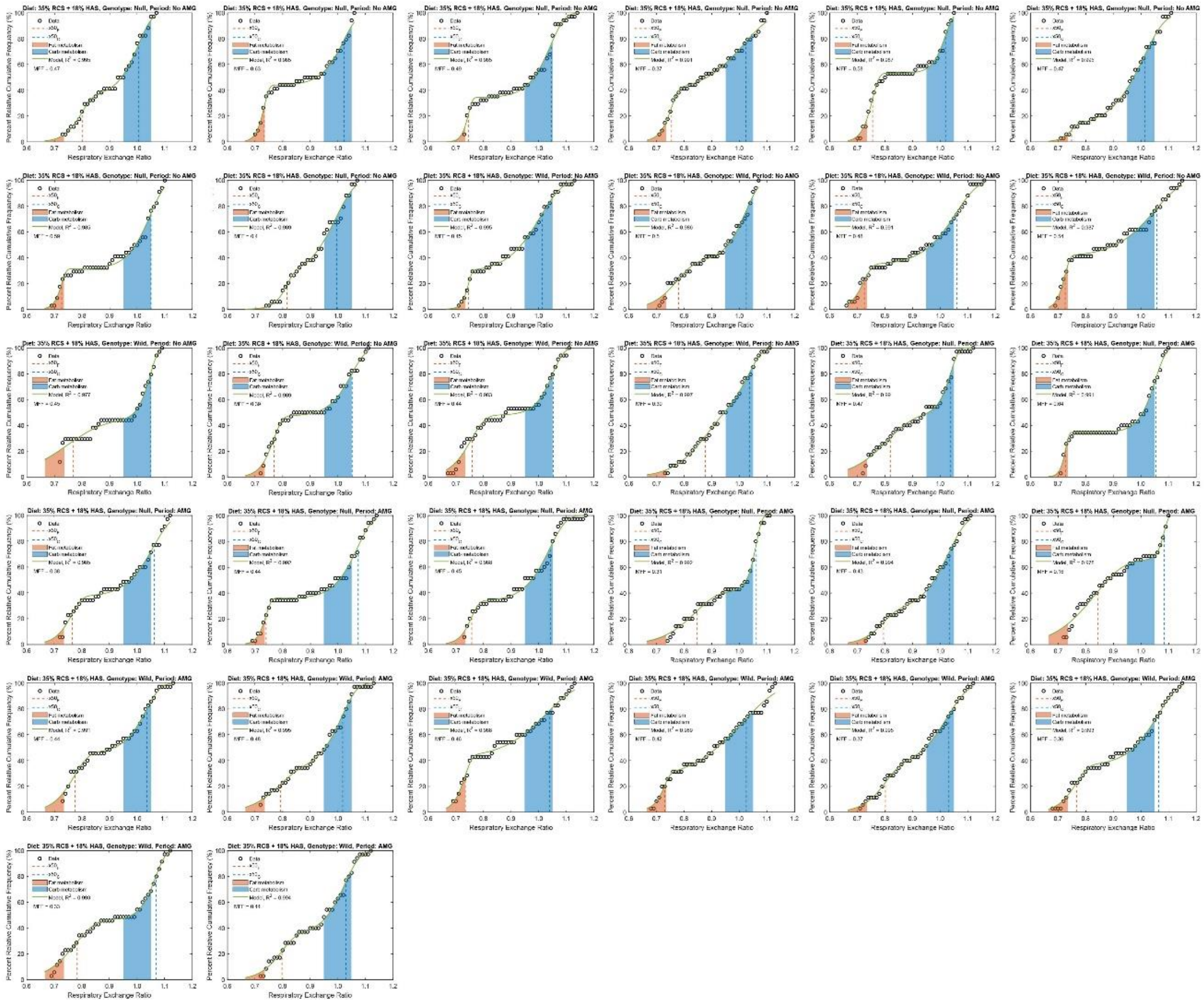

**Fig. S5.** Data and curve fits for Percent Relative Cumulative Frequency (PRCF) modeling using the Mixed Weibull Cumulative Distribution function for all individual mice on the Intermediate SDS diet. Red shading represents the 'ideal' fat oxidation range of respiratory exchange ratio (RER), while the blue shading represents the 'ideal' carbohydrate range of RER values.

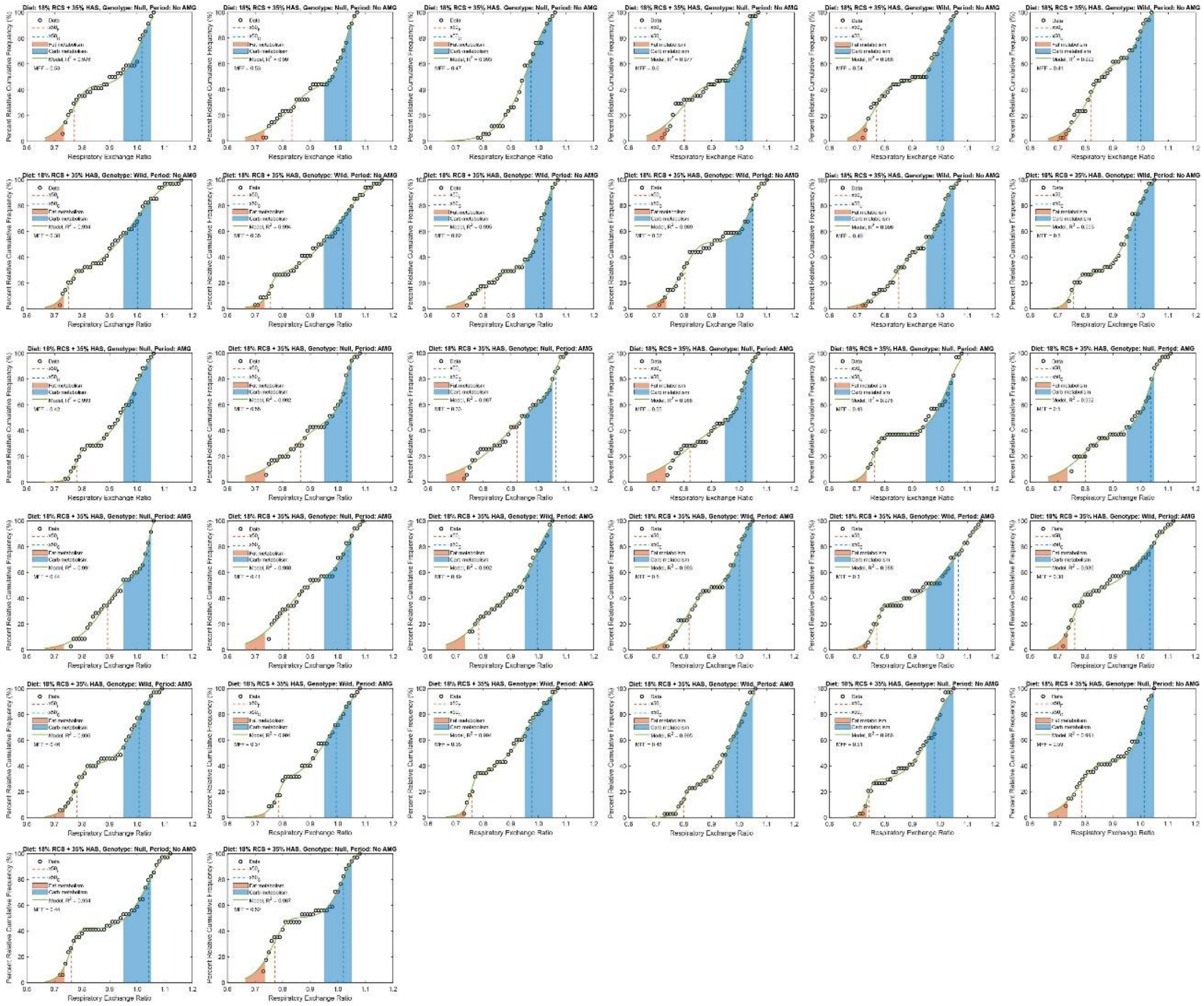

**Fig. S6.** Data and curve fits for Percent Relative Cumulative Frequency (PRCF) modeling using the Mixed Weibull Cumulative Distribution function for all individual mice on the Intermediate RS diet. Red shading represents the 'ideal' fat oxidation range of respiratory exchange ratio (RER), while the blue shading represents the 'ideal' carbohydrate range of RER values.

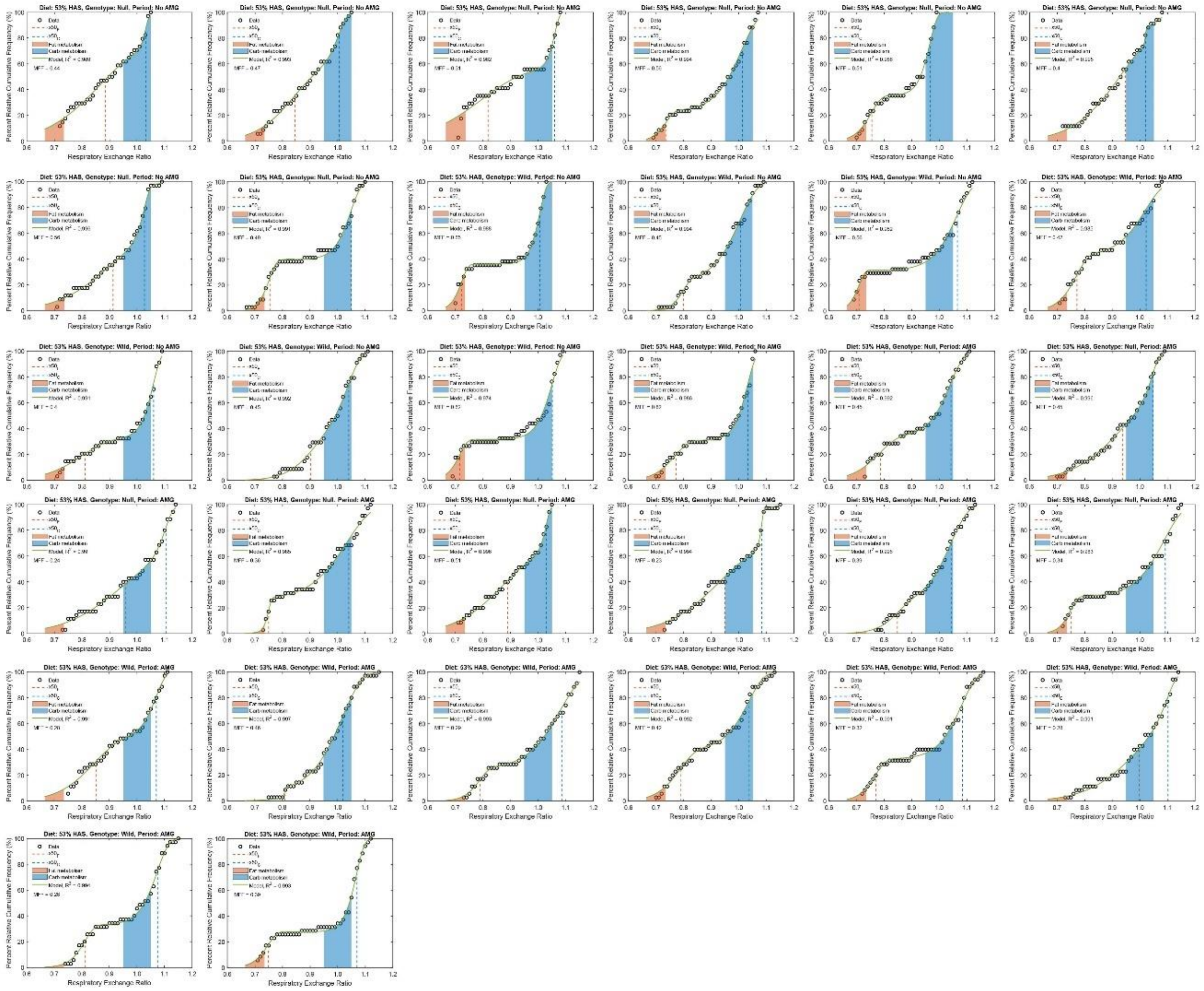

**Fig. S7.** Data and curve fits for Percent Relative Cumulative Frequency (PRCF) modeling using the Mixed Weibull Cumulative Distribution function for all individual mice on the High RS diet. Red shading represents the ‘ideal’ fat oxidation range of respiratory exchange ratio (RER), while the blue shading represents the ‘ideal’ carbohydrate range of RER values.

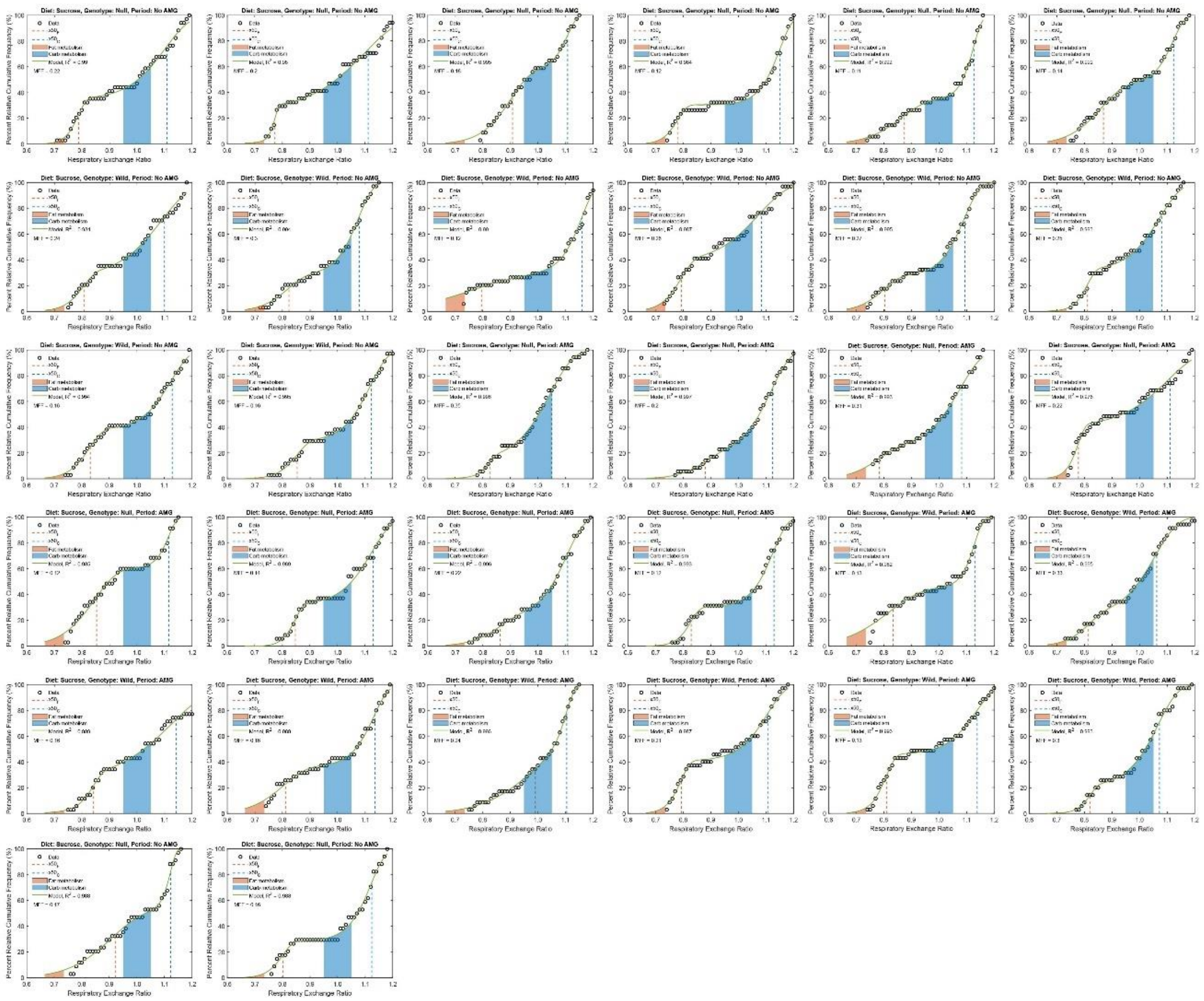

**Fig. S8.** Data and curve fits for Percent Relative Cumulative Frequency (PRCF) modeling using the Mixed Weibull Cumulative Distribution function for all individual mice on the Sucrose diet. Red shading represents the ‘ideal’ fat oxidation range of respiratory exchange ratio (RER), while the blue shading represents the ‘ideal’ carbohydrate range of RER values.

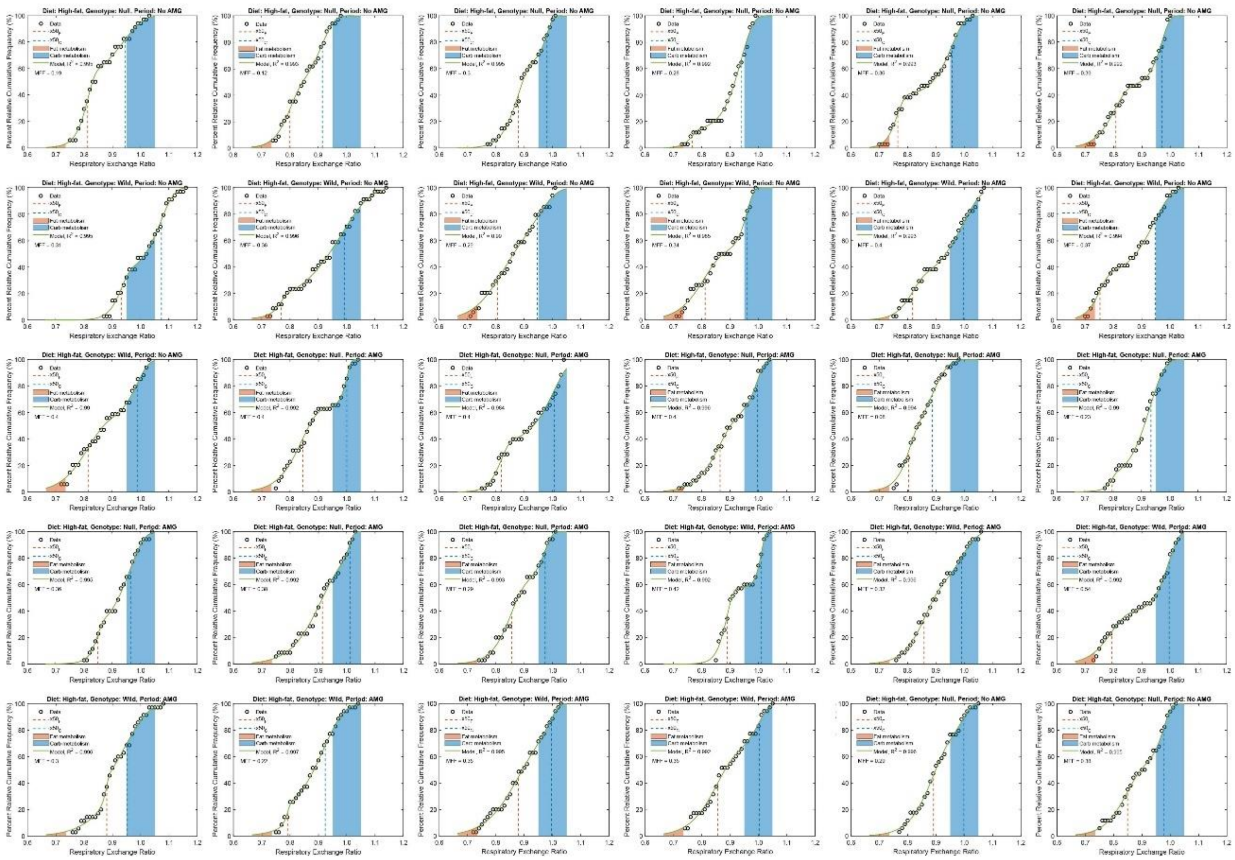

**Fig. S9.** Data and curve fits for Percent Relative Cumulative Frequency (PRCF) modeling using the Mixed Weibull Cumulative Distribution function for all individual mice on the High-fat diet. Red shading represents the ‘ideal’ fat oxidation range of respiratory exchange ratio (RER), while the blue shading represents the ‘ideal’ carbohydrate range of RER values.

**Table S1.** Experimental diet compositions.

| Component | High SDS TD.01629 | Intermediate SDS TD.02130 | Intermediate RS TD.02130 | High RS TD.02130 | Sucrose TD.02129 | High-fat TD.88137 |
|---|---|---|---|---|---|---|
| | | | g/kg | | | |
| Casein | 200 | 200 | 200 | 200 | 200 | 195 |
| DL-Methionine | 3 | 3 | 3 | 3 | 3 | 3 |
| Maltodextrin | 120 | 120 | 120 | 120 | 0 | 0 |
| Raw corn starch | 530 | 353 | 177 | 0 | 0 | 150 |
| Resistant starch (Novelose 260) | 0 | 177 | 353 | 530 | 0 | 0 |
| Sucrose | 0 | 0 | 0 | 0 | 650 | 342 |
| Anhydrous milkfat | 0 | 0 | 0 | 0 | 0 | 210 |
| Cholesterol | 0 | 0 | 0 | 0 | 0 | 1.5 |
| Soybean Oil | 50 | 50 | 50 | 50 | 50 | 0 |
| Cellulose | 50 | 50 | 50 | 50 | 50 | 50 |
| Mineral mix[a] | 35 | 35 | 35 | 35 | 35 | 35 |
| Calcium carbonate | 0 | 0 | 0 | 0 | 0 | 4 |
| Vitamin mix[b] | 10 | 10 | 10 | 10 | 10 | 10 |
| Choline bitartrate | 2.5 | 2.5 | 2.5 | 2.5 | 2.5 | 0 |
| TBHQ (antioxidant) | 0.01 | 0.01 | 0.01 | 0.01 | 0.01 | 0 |
| Ethoxyquin | 0 | 0 | 0 | 0 | 0 | 0.04 |
| | | | % kcal | | | |
| Protein | 19.6 | 19.6 | 19.6 | 19.6 | 19.6 | 15.2 |
| Carbohydrate | 67.4 | 67.4 | 67.4 | 67.4 | 67.4 | 42.7 |
| Fat | 13.0 | 13.0 | 13.0 | 13.0 | 13.0 | 42.0 |
| | | | % by weight | | | |
| Protein | 17.7 | 17.7 | 17.7 | 17.7 | 17.7 | 17.3 |
| Carbohydrate | 60.9[c] | 60.9[c] | 60.9 | 60.9 | 60.9 | 48.5 |
| Fat | 5.2 | 5.2 | 5.2 | 5.2 | 5.2 | 21.2 |
| Energy density (kcal/g) | 3.6 | 3.6 | 3.6 | 3.6 | 3.6 | 4.5 |

AMG, amyloglucosidase; RER, respiratory exchange ratio.

[a]Mineral mix composition (g/kg; AIN-93G-MX, TD.94046): calcium carbonate, 357.0; potassium phosphate (monobasic), 196.0; potassium citrate (monohydrate), 70.78; sodium chloride, 74.0;

potassium sulfate, 46.6; magnesium oxide, 24.3; ferric citrate, 6.06; zinc carbonate, 1.65; manganous carbonate, 0.63; cupric carbonate, 0.31; potassium iodate, 0.01; sodium selenate, 0.0103; ammonium paramolybdate (tetrahydrate), 0.008; sodium meta-silicate (nonahydrate), 1.45; chromium potassium sulfate (dodecahydrate), 0.275; lithium chloride, 0.0174; boric acid, 0.0815; sodium fluoride, 0.0635; nickel carbonate hydroxide (tetrahydrate), 0.0318; ammonium meta-vanadate, 0.0066; sucrose (fine ground), 220.7. Note that these values are the g/kg amounts within the mineral mix, and only 35 g/kg of this mix was used within the diet.

[b]Vitamin mix composition (g/kg; AIN-93-VX, TD.94047): niacin, 3.0; calcium pantothenate, 1.6; pyridoxine hydrochloric acid, 0.7; thiamin (81%), 0.6; riboflavin, 0.6; folic acid, 0.2; biotin, 0.02; vitamin $B_{12}$ (0.1% in mannitol), 2.5; vitamin E (DL-alpha tocopheryl acetate, 500 IU/g), 15.0; vitamin A palmitate (500,000 IU/g), 0.8; vitamin $D_3$ (cholecalciferol, 500,000 IU/g), 0.2; vitamin $K_1$ (phylloquinone), 0.075; sucrose (fine ground), 974.7. Note that these values are the g/kg amounts within the mineral mix, and only 10 g/kg of this mix was used within the diet.

[c]53% of which was experimental carbohydrate (High SDS [corn starch], High RS [Novelose 260]).

**Table S2.** Contents of rapidly digestible starch (RDS), slowly digestible starch (SDS), and resistant starch (RS) in different diets (%, dry starch basis).[a] Mean values shown with ± standard deviation as applicable.

| Diet | Rapidly digestible starch (%) | Slowly digestible starch (%) | Resistant starch (%) | Percent amylose[b] (%) | Food quotient[c] |
|---|---|---|---|---|---|
| High SDS diet | 74.0 ± 4.8 | 26.9 ± 5.4 | -0.9 | 23 | 0.93 |
| Intermediate SDS diet | 64.7 ± 2.7 | 25.1 ± 2.4 | 10.2 | 34 | 0.93 |
| Intermediate RS diet | 59.5 ± 3.7 | 25.2 ± 6.5 | 15.3 | 46 | 0.93 |
| High RS diet | 55.1 ± 2.8 | 16.4 ± 3.0 | 28.5 | 57 | 0.93 |
| Sucrose diet[d] | 32.2[b] | 32.7[b] | 0.0 | 0 | 0.93 |
| High-fat diet | n.d. | n.d. | n.d. | 9 | 0.85 |
| PicoLab diet 5053 (non-experimental) | 58.7 ± 4.7 | 43.7 ± 3.1 | -2.4 | 18 | 0.92 |

AMG, amyloglucosidase; n.d., not determined; RS, resistant starch; SDS, slowly digestible starch.

[a]Analyzed using the Englyst assay (K. N. Englyst et al. 1999; H. N. Englyst, Kingman, and Cummings 1992).

[b]Calculated as percent of amylose within the digestible carbohydrate component of each diet, considering normal corn starch contains 28% amylose and Novelose 260 contains 70% amylose.

[c]Food quotient of oxidation per diet: calculated according to macronutrient ratio by weight, assuming a quotient of oxidation of 1.0 for carbohydrates, 0.70 for fats, and 0.825 for proteins.
[d]Glucose contents of the sample incubated after 20 and 120 min were 32.2 and 32.7%, respectively. This is due to the invertase in the Englyst assay, noting that the other carbohydrate components of this diet are fructose and cellulose and thus would not be detected using the Englyst assay.

#### Supplementary Information References
